# Loss of restriction-modification methyltransferases drives persistence to fluoroquinolones in *Pseudomonas aeruginosa*

**DOI:** 10.64898/2026.09.26.754597

**Authors:** Maureen Micaletto, Nicolas Oswaldo Gomez, Anna Olina, Aleksei Agapov, Urszula Łapińska, Matthias Preuße, María García-Castillo, Christian Hacker, Rafael Canton, Stineke van Houte, Susanne Häussler, Edze R. Westra, Multi-Defence Consortium, ERADIAMR Consortium, Stefano Pagliara

## Abstract

The resurgence of phage therapy has renewed interest in the interplay between phage resistance and antibiotic susceptibility^1,2,3^ and has stimulated research in the emergent field of non-canonical cellular functions carried out by defence systems^4^. Yet it remains largely unknown whether intracellular antiphage defence systems influence bacterial physiology, resistance or persistence to antibiotics. Here we discovered that besides its canonical antiphage defence function, the type I restriction modification (RM) system profoundly affects the physiology of the opportunistic pathogen *Pseudomonas aeruginosa*. Deletion of the type I RM methyltransferase HsdM reduces the size of the bacterial nucleoid and delays DNA replication initiation and exit from lag phase in *P. aeruginosa* PAO1. Crucially, *P. aeruginosa* strains isolated from patients with cystic fibrosis (CF) and lacking the RM type I system also display slower growth compared to strains isolated from other sites of infections and encoding this system. Deletion of HsdM selectively increases the levels of persisters that survive treatment with fluoroquinolones by displaying enhanced SOS response but without acquiring resistance. Importantly, we measured elevated persistence to fluoroquinolones also in *P. aeruginosa* CF isolates lacking the type I RM system, providing a functional link between RM systems, slow growth and persistence to fluoroquinolones. Together these findings open a new way of approaching bacterial susceptibility to antibiotics, bringing antiphage defence systems in a forward-facing position in this field.

## Introduction

Antibiotic treatment failure is a growing global health threat^5^. Although genetic resistance is a major contributor to treatment failure, many chronic and recurrent infections persist in the absence of heritable resistance mutations due to the presence of subpopulations that survive otherwise lethal antibiotic exposure^6,7,8,9^. These persister cells are in a transient physiological state typically characterized by reduced growth rates^8,9^, elevated SOS response and DNA damage repair^10,11^, reduced antibiotic accumulation and intracellular pH^12,13^, and increased toxin-to-antitoxin ratios^6^.

Since resistance and persistence to antibiotics often complicate treatment^14,15^, research is ongoing to treat chronic and recurrent bacterial infections with bacteriophages^16,17^ alongside antibiotics and to understand the relationship between phage resistance and antibiotic resistance^18,19^. In several cases, resistance to phages generates collateral sensitivity to antibiotics, creating exploitable therapeutic trade-off^3,18,19^. However, such effects have been studied almost exclusively in the context of phage resistance mediated by alterations to cell-surface receptors^20,21^.

Recent genomic surveys have revealed a vast repertoire of anti-phage defence systems, comprising hundreds of distinct mechanisms that collectively rival the complexity of immune systems in eukaryotes^22,23,24^. Although these systems are best known for protecting bacteria from phage infection, emerging evidence suggests that they can influence fundamental aspects of cellular physiology^4,25,26^. Moreover, a recent study showed that the CBASS antiphage system can increase the susceptibility of *Vibrio cholerae* to treatment with antifolate antibiotics^2^. Crucially, whether this plethora of intracellular defence systems influence antibiotic resistance or persistence more broadly remains largely unknown. *Pseudomonas aeruginosa* is an excellent model to investigate this hypothesis considering that cystic fibrosis patients experiencing chronic *P. aeruginosa* lung infections are increasingly being treated with phage-antibiotic therapy^27^ and that bioinformatic analysis showed reduced prevalence of antiphage defence systems in *P. aeruginosa* strains from cystic fibrosis patients^28^.

Here we systematically investigate the impact of known defence systems on persistence to antibiotics using a derivative of *P. aeruginosa* PAO1 lacking all known antiphage defence systems^29^. We identify a type I restriction–modification (RM) system as a major determinant of persistence to DNA-targeting antibiotics. We employed transcriptomics, electron microscopy, biochemical assays and microfluidics-based single-cell analysis to understand how the loss of the type I RM methyltransferase affects the growth and persistence of *P. aeruginosa*. Crucially, we show that slow-growing cystic fibrosis isolates lacking RM systems also display elevated persistence to DNA-targeting antibiotics, therefore offering a new way of tackling persistence to antibiotics.

## Results

### The loss of the type I restriction–modification system selectively enhances persistence to DNA-targeting antibiotics

To test whether bacterial immune systems influence antibiotic resistance or persistence, we compared wild-type *Pseudomonas aeruginosa* PAO1 with a derivative lacking all three chromosomally encoded defence systems, i.e. Helicase–DUF2290, Gabija and a type I restriction–modification (RM) system, as well as the prophages Pf1 and phiCTX (defenceless PAO1; dPAO1)^29^. We employed bactericidal antibiotics targeting translation (tobramycin), cell-wall synthesis (meropenem and ceftazidime), membrane integrity (polymyxin B) and DNA replication (ciprofloxacin).

Minimum inhibitory concentration (MIC) values were not different between PAO1 and dPAO1 for all antibiotics tested (Extended Data Fig. 1a), indicating that deletion of these defence systems does not alter population-level susceptibility. Time-kill assays performed at 10× the MIC value of each antibiotic for 6 hours resolved biphasic killing dynamics characteristic of persister formation (Extended Data Fig. 1b-e). Regrowth of survivors in antibiotic-free medium followed by re-exposure to the same antibiotic yielded the original survival fraction and biphasic dynamics, excluding stable resistance and confirming a reversible, non-heritable phenotype^6,30^. Deletion of defence systems had no effect on persister levels to tobramycin, meropenem, ceftazidime or polymyxin B. In contrast, dPAO1 exhibited a >2-log increase in persisters to the fluoroquinolone ciprofloxacin relative to PAO1 (Fig. 1a and Extended Data Fig. 1e). A similar increase in survival was observed with the fluoroquinolones ofloxacin, moxifloxacin and levofloxacin (Extended Data Fig. 1f), indicating selective persistence to DNA-targeting antibiotics.

**Figure 1.**
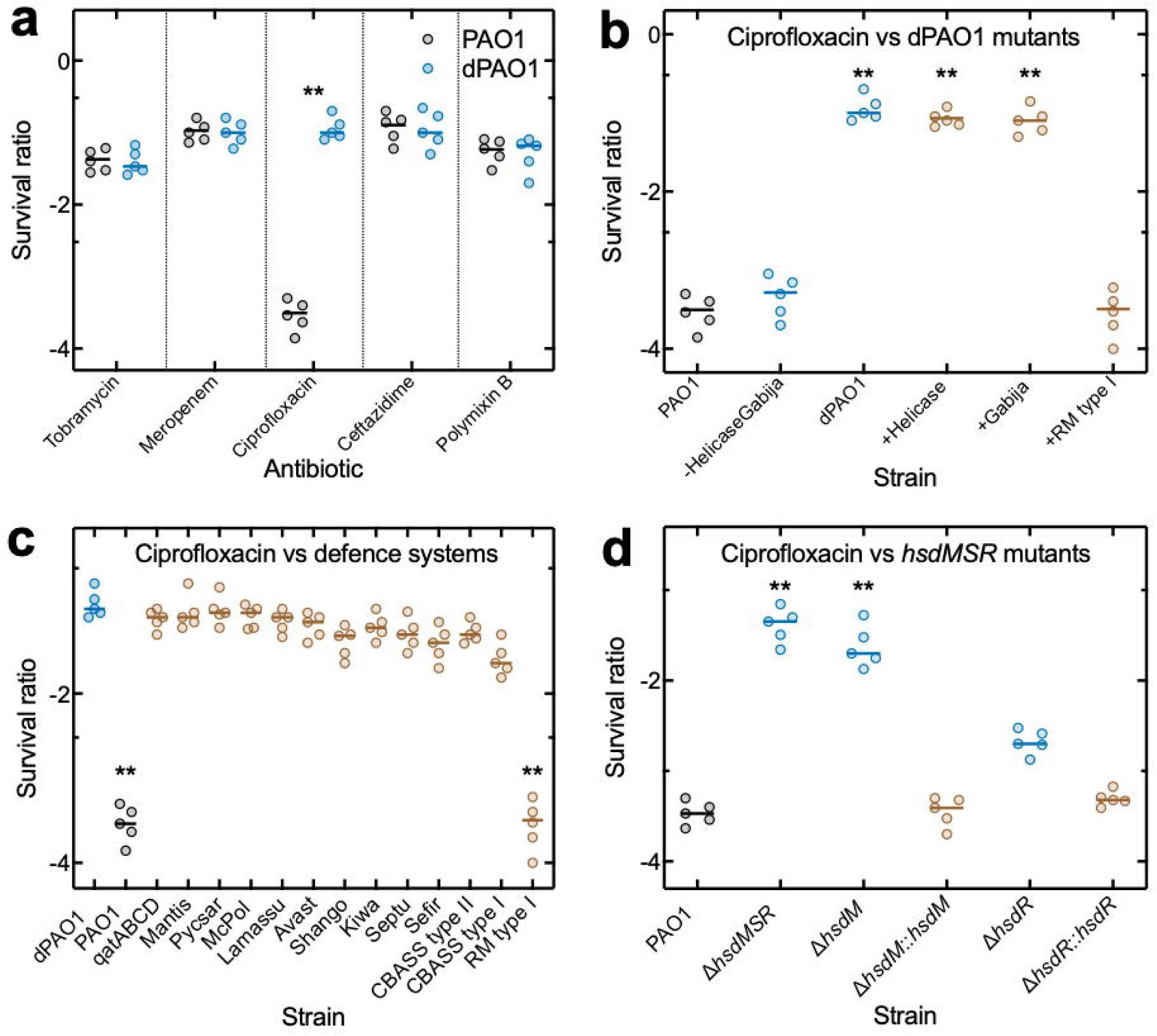
The type I RM methyltransferase limits persistence to ciprofloxacin in *Pseudomonas aeruginosa*. **a**, Survival ratio of the *P. aeruginosa* PAO1 strain (grey circles) and the isogenic derivative strain dPAO1 (blue circles) after 4 h exposure to bactericidal antibiotics at 10× their MIC value and regrowth on plates in antibiotic-free medium. **b**, Survival ratio of PAO1, PAO1 with all systems deleted apart from RM type I, dPAO1 and dPAO1 with individual deleted defence systems reintroduced via the pLOLA plasmid after 4 h exposure to ciprofloxacin at 10× its MIC value and regrowth on plates. **c**, Survival ratio of PAO1, dPAO1 and dPAO1 in which heterologous defence modules were introduced via the pLOLA plasmid after 4 h exposure to ciprofloxacin at 10× its MIC value and regrowth on plates. **d**, Survival ratio of the PAO1, Δ*hsdMSR*, Δ*hsdM*, Δ*hsdR* Δ*hsd*M*::hsdM* and Δ*hsdR::hsdR* strains after 4 h exposure to ciprofloxacin at 10× its MIC value and regrowth on plates. The survival ratio has been calculated as the ratio of colony forming units after each treatment compared to the corresponding colony forming units before treatment and the log_10_ of these values has been plotted in the graphs. Each point represents an independent biological replicate (n=5). The centre line indicates the median. Statistical significance was determined using two-tailed Mann–Whitney U tests for pairwise comparisons vs the PAO1 strain with the exception of panel c where tests were performed vs the dPAO1 strain; ** indicate a p-value < 0.01.

Retaining the type I RM system as the sole chromosomal defence locus or restoring this system in dPAO1 via the pLOLA expression vector^31^ fully reduced persister levels (Fig. 1b), demonstrating that this defence system constrains persistence to DNA-targeting antibiotics, besides its canonical protection against a wide range of phylogenetically diverse lytic and temperate phages (Extended Data Fig. 2).

We next asked whether the introduction of other known defence systems could increase persistence. Twelve heterologous defence systems absent from PAO1— including representatives of CBASS, Lamassu, Shango and Kiwa—were introduced into dPAO1 via the pLOLA expression vector^31^. None altered ciprofloxacin persistence under the conditions tested (Fig. 1c), indicating that elevated persistence is a specific feature of the endogenous type I RM system.

The PAO1 type I RM locus comprises *hsdM* (methyltransferase), *hsdS* (sequence specificity) and *hsdR* (restriction endonuclease). Deletion of the entire locus (Δ*hsdMSR*) reproduced the high persistent phenotype observed in dPAO1. Strikingly, deletion of *hsdM* alone fully phenocopied this persistent state, whereas deletion of *hsdR* produced only a modest increase in survival (Fig. 1d). Chromosomal complementation of *hsdM* or *hsdR* restored low wild-type persistence levels (Fig. 1d). Because HsdM and HsdS form the M₂S₁ methyltransferase complex required for assembly of the active R₂M₂S₁ restriction holoenzyme^32^, loss of HsdM abrogates coordinated RM activity (Extended Data Fig. 3). These results demonstrate that disruption of RM-dependent methylation—rather than restriction cleavage per se— drives high persistence to fluoroquinolones.

Together, these results identify the type I RM methyltransferase in *P. aeruginosa*PAO1 as a specific regulator of persistence to DNA-targeting antibiotics.

### The loss of type I RM methyltransferase delays single-cell growth resumption and increases the frequency of persisters that survive ciprofloxacin

Next, we employed a mother-machine microfluidics platform adapted for *P. aeruginosa*^12^, to investigate how the type I RM methyltransferase shapes cellular heterogeneity during exposure to the fluoroquinolone ciprofloxacin. Following 4 h exposure to ciprofloxacin at 10× its MIC value and subsequent recovery in antibiotic-free medium, we identified persisters that remained non-dividing during antibiotic treatment and resumed growth upon exposure to antibiotic-free medium, fulfilling the canonical definition of persistence^6,30^ (Fig. 2a,b). Persisters levels were around 0.5% in the dPAO1, Δ*hsdMSR* and Δ*hsdM* strains, whereas were not detected in wild-type PAO1 or Δ*hsdR* strains (Fig. 2c). Besides this striking difference, the dPAO1, Δ*hsdMSR* and Δ*hsdM* strains also displayed significantly lower levels of cells that filamented or lysed and higher levels of cells that underwent limited cell doubling and then stopped growing (Fig. 2a-c). Taken together these data demonstrate that the loss of the type I RM methyltransferase shifts the population structure towards survival-associated phenotypes against DNA-damaging antibiotics, enhancing the probability that individual cells enter a protected, non-dividing state during antibiotic stress.

**Fig. 2.**
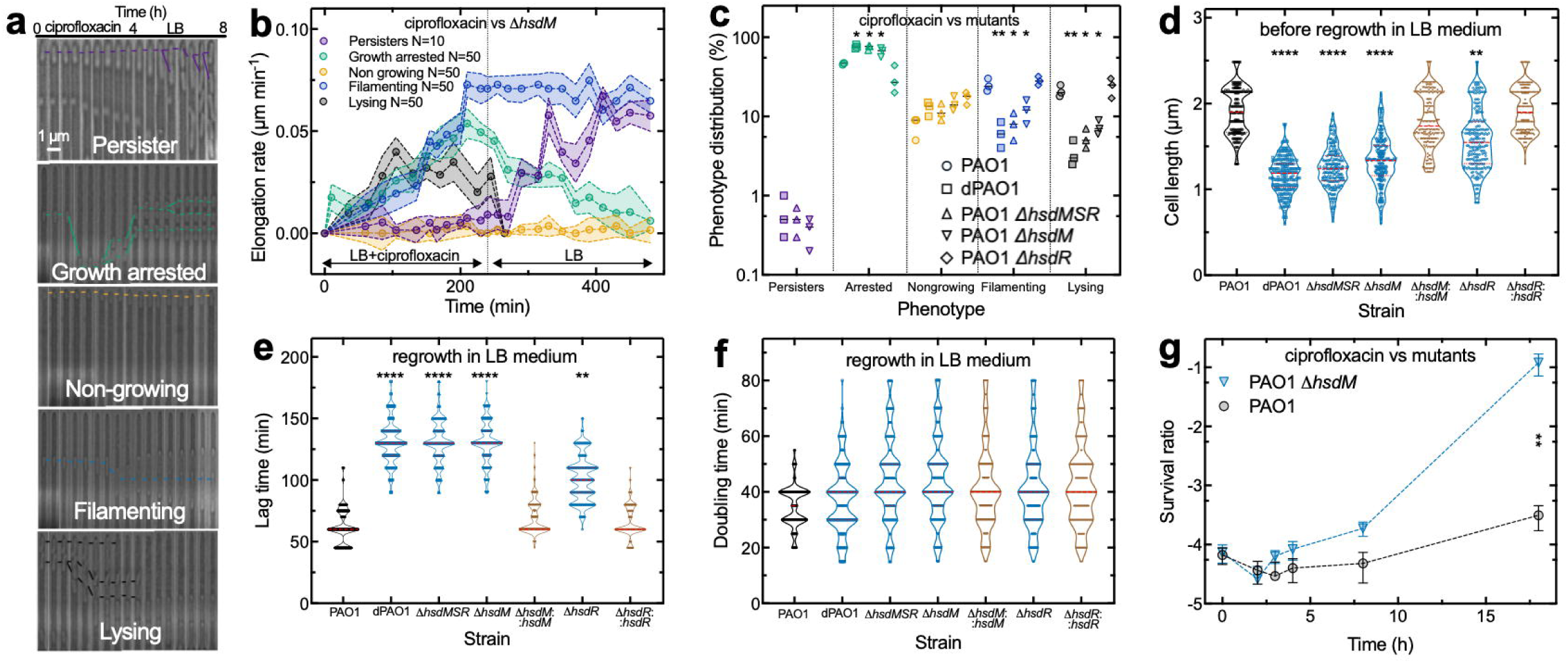
The loss of type I RM methyltransferase delays single-cell growth resumption and increases single-cell survival to ciprofloxacin. a, Representative mother-machine time-lapse images showing five distinct single-cell fates during 4h exposure of the PAO1 Δ*hsdM* strain to ciprofloxacin at 10× its MIC value followed by antibiotic removal and incubation in LB medium: persisters (purple lines), growth arrested cells (green), non-growing cells (yellow), filamenting cells (blue) and lysing cells (black). **b,** Corresponding average elongation rates over time stratified by phenotype. Points and shaded areas indicate the mean and the standard deviation of elongation rates for n=50 individual cells per phenotypes, except for persister cells with n=10. **c,** Corresponding quantification of cell-fate distributions for the PAO1 (circles), dPAO1 (squares), Δ*hsdMSR* (upward triangles), Δ*hsdM* (downward triangles) and Δ*hsdR* (diamonds) strains. Statistical significance was determined using lognormal Welch t-test; *: p-value < 0.05; **: p-value < 0.01. **d,** Distribution of the length of individual cells when loaded in the mother machine device from stationary phase cultures. n=231, 224, 225, 246, 231, 247 and 224 for the PAO1, dPAO1, Δ*hsdMSR*, Δ*hsdM*, Δ*hsdR* Δ*hsd*M*::hsdM* Δ*hsdR::hsdR* strains, respectively. **e,** Distribution of lag times of individual cells during growth in antibiotic-free medium. n=225, 227, 226, 226, 220, 223 and 223 for the PAO1, dPAO1, Δ*hsdMSR*, Δ*hsdM*, Δ*hsdR* Δ*hsd*M*::hsdM* Δ*hsdR::hsdR* strains, respectively. **f,** Corresponding distribution of doubling times of individual cells during growth in antibiotic-free medium. n=229, 231, 237, 240, 228, 229 and 277 for the PAO1, dPAO1, Δ*hsdMSR*, Δ*hsdM*, Δ*hsdR* Δ*hsd*M*::hsdM* Δ*hsdR::hsdR* strains, respectively. Red dashed lines in d-f indicate the median and quartiles of each disrtibution. Statistical significance was determined using two-tailed Mann–Whitney U tests vs the PAO1 strain; **: p-value < 0.01; ****: p-value < 0.0001. **g,** Temporal dynamics of survival ratio of growing PAO1 and Δ*hsdM* cultures exposed at the indicated time points to ciprofloxacin at 10× its MIC value and regrowth on antibiotic-free plates. Points represent the mean and standard deviation of three independent biological replicates; **: p-value < 0.01 (two-tailed unpaired t-test).

Next, we tested whether the type I RM methyltransferase has an impact on bacterial growth even in the absence of antibiotic treatment. The dPAO1, Δ*hsdMSR* and Δ*hsdM* strains displayed reduced cell length in stationary phase (Fig. 2d, Extended Data Table 1) and a prolonged lag phase (Fig. 2e; Extended Data Table 2), but similar doubling times during exponential growth (Fig. 2f; Extended Data Table 3) compared to the PAO1, Δ*hsdR* or complemented strains. Thus, disruption of the type I RM methyltransferase establishes a profoundly altered growth resumption besides affecting survival to fluoroquinolones. Consistent with this growth resumption-specific phenotype, disruption of *hsdM* led to increased persistence during regrowth from stationary phase but did not have an impact on persister levels during exponential growth (Fig. 2g).

Together, these observations reveal a previously unknown function of the type I RM methyltransferase that is the regulation of single-cell growth dynamics and survival to DNA damaging antibiotics.

### The loss of type I RM methyltransferase triggers a shrinking of the nucleoid, a delay in chromosome replication and an increase in the SOS response

The PAO1 type I RM system has been previously linked to gene regulation^33^. We therefore compared the transcriptomes of the Δ*hsdM* mutant and the wild-type PAO1 strain under two conditions: (i) after 17 h of growth and (ii) during early regrowth from stationary phase (lag phase), when increased persistence to fluoroquinolones is observed.

During stationary phase, the Δ*hsdM* and wild-type strain showed minimal transcriptional differences, restricted to the deleted *hsdM* locus, its cognate restriction enzyme *hsdR*, and a single upregulated universal stress protein (Extended Data Fig. 4a). In contrast, during lag phase, Δ*hsdM* cells exhibited a defined transcriptional shift (Fig. 3a, Extended Data Fig. 4b). Functional enrichment revealed a rewiring of glycolysis, pyruvate metabolism and nucleotide biosynthesis alongside glycolysis, pyruvate metabolism and nucleotide biosynthesis, alongside coordinated activation of two-component signaling, stress response, and transport processes, together with modules controlling iron, heme, metal, and redox homeostasis, indicative of engagement of protective, redox-sensitive pathways (Fig. 3b).

**Fig. 3.**
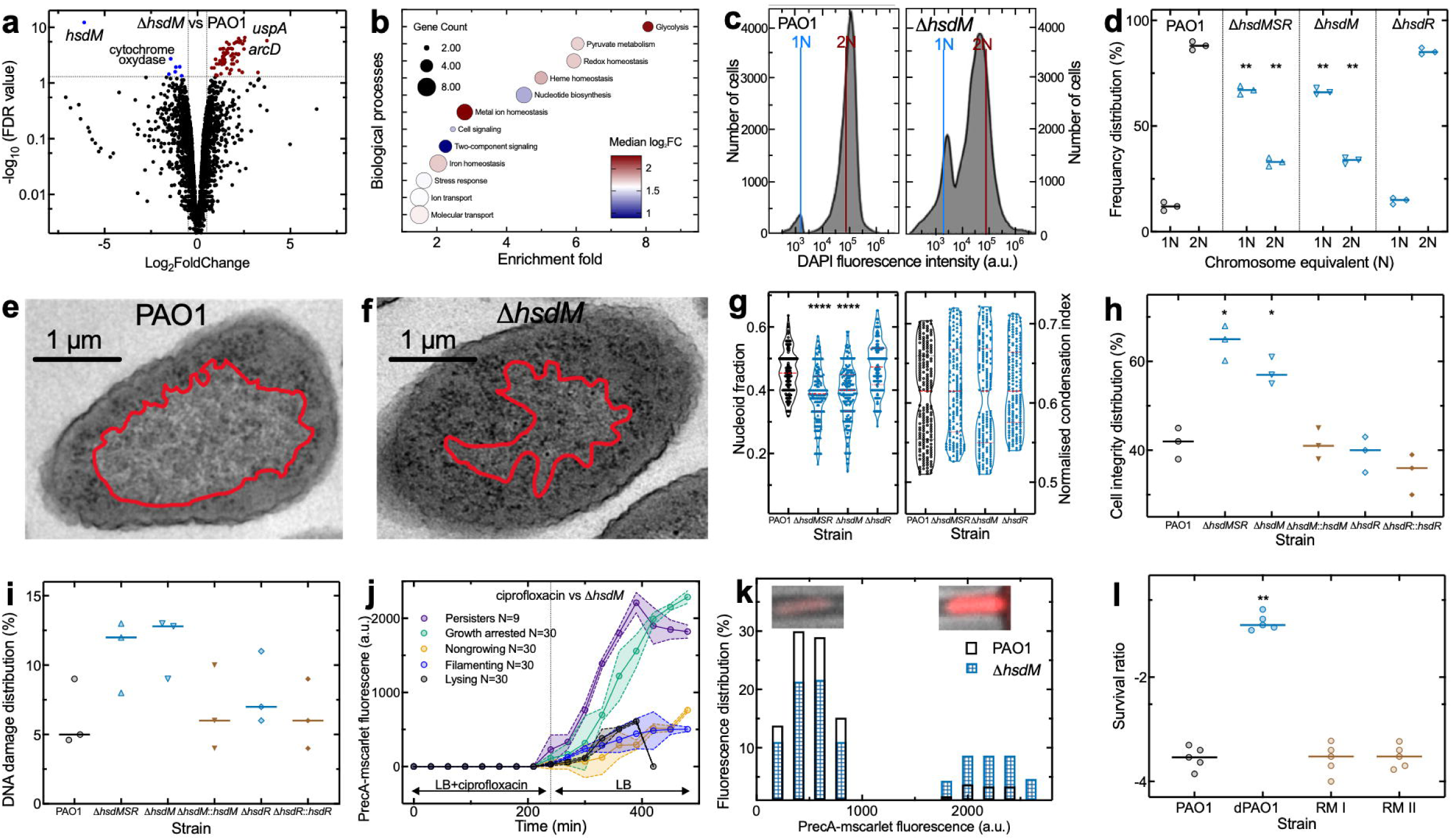
The loss of type I RM methyltransferase triggers a reduction of the nucleoid size, a delay in chromosome replication and an increase in SOS response. a, Volcano plot of differential gene expression between the Δ*hsdM* and the PAO1 parental strain during lag phase. Red and blue points indicate significantly up-and down-regulated genes, respectively. **b**, Corresponding functional enrichment analysis of differentially expressed genes revealing enriched biological processes presented as circles of different size (indicating gene counts) and colour (representing median log₂ fold change). **c,** Number of chromosome equivalents (*N*) in individual cells of the PAO1 (left) and Δ*hsdM* (right) strains in lag phase after 120 min of incubation with rifampicin (200μg/mL) and aztreonam (10μg/mL). Corresponding 1*N* stationary phase cells were used as a reference. The results are representative of three independent experiments consisting of n=50,000 individual bacteria each. **d,** Corresponding percentage of cells containing one (1N) or two (2N) chromosome equivalents in the PAO1, Δ*hsdMSR*, Δ*hsdM* and Δ*hsdR* strains. Each point represents an independent biological replicate. The centre line indicates the median. Statistical significance was determined using Mann-Withney test vs the PAO1 strain, **: p-value < 0.01. **e-f,** Representative transmission electron micrographs of the PAO1 and Δ*hsdM* (right) bacteria with their nucleoids outlined in red. **g,** Corresponding distribution of nucleoid occupancy relative to total cell area (left) and of Fourier transform–based normalised condensation index of nucleoids in the PAO1, Δ*hsdMSR*, Δ*hsdM* and Δ*hsdR* strains. Statistical significance was determined using two-tailed Mann–Whitney U tests vs the PAO1 strain. n=159, 181,168, and 159 for the PAO1, Δ*hsdM*, Δ*hsdR* and Δ*hsdMSR* respectively. ****: p-value < 0.0001. **h,** Percentage of bacteria with intact cell integrity after treatment with ciprofloxacin at 10× its MIC value. Cell integrity was evaluated by forward scatter (FSC) and side scatter (SSC) parameters, which reflect cell size and granularity, respectively. A gate was established based on the FSC–SSC profile of the non-treated overnight culture to exclude debris and dead cells (see Extended Data Figure 4e). Only events falling within this gate were included in the intact cells count. Each point represents an independent biological replicate consisting of n=50,000 individual bacteria each. The centre line indicates the median. Statistical significance was determined using two-tailed Mann–Whitney U tests vs the PAO1 strain. *: p-value < 0.05. **i,** Percentage of cells displaying DNA strand breaks after exposure to ciprofloxacin at 10× its MIC value. n=50,000 cells for each strain. Each point represents an independent biological replicate consisting of n=50,000 individual bacteria each. The centre line indicates the median. **j** Temporal dynamics of P*recA-mscarlett* fluorescence during and after treatment with ciprofloxacin at 10× its MIC stratified by cell fate for the Δ*hsdM* strain. Points and shaded areas represent the mean and standard deviation of n=30 individual cells per phenotype with n=10 for persisters. **k,** Corresponding single-cell distribution of P*recA-mscarlett* fluorescence intensity at the last time point for the PAO1 (open bars) and the Δ*hsdM* strain (checked bars). Insets: representative fluorescence microscopy images of low-and high-fluorescence cells of the Δ*hsdM* strain. **l,** Survival ratio of PAO1, dPAO1 and dPAO1 complemented with pLOLA plasmid encoded type I or II RM systems after exposure to ciprofloxacin at 10× its MIC value. Each point represents an independent biological replicate, the centre line indicates the median. Statistical significance was determined using two-tailed Mann–Whitney U tests vs the PAO1 strain. **: p-value < 0.01.

Given the specificity of persistence to DNA-targeting antibiotics, we next examined chromosome replication during lag phase via DAPI staining^34^. The Δ*hsdMSR* and Δ*hsdM* strains displayed significantly less cells with two chromosome equivalents and broader DAPI fluorescence distributions compared to the wild-type and the Δ*hsdR* strains (Fig. 3c,d Extended Data Fig. 4c), indicating delayed replication initiation and increased replication-state heterogeneity in the absence of antibiotic treatment.

We then assessed chromosome organization by transmission electron microscopy (Fig. 3e). The Δ*hsdMSR* and Δ*hsdM* strains exhibited reduced nucleoid occupancy with DNA confined to a smaller intracellular region compared to the wild-type and the Δ*hsdR* strains, while Fourier transform–based analysis revealed no change in local DNA packing density (Fig. 3g, Extended Data Fig. 4d). These data indicate that the type I RM methyltransferase modulates large-scale chromosome organization.

Next, we determined the impact of the loss of *hsdM* on cellular and DNA damage caused by ciprofloxacin, using a terminal deoxynucleotidyl transferase dUTP nick end-labelling (TUNEL) assay^35^ and flow cytometry. Following exposure to a supra-inhibitory concentration of ciprofloxacin (10× MIC), the Δ*hsdMSR* and Δ*hsdM* strains retained significantly higher proportions of structurally intact cells than the wild-type and Δ*hsdR* strains (Fig. 3h), in which extensive cell lysis occurred (Extended Data Fig. 4e), however all strains exhibited similarly low levels of detectable double-strand breaks (Fig. 3i, Extended Data Fig 4f). Moreover, the Δ*hsdMSR* and Δ*hsdM* strains accumulated significantly more DNA damage than the wild type and Δ*hsdR* strains after exposure to ciprofloxacin at its MIC value (Extended Data Fig. 4g). Therefore, increased persistence to fluoroquinolones in the absence of *hsdM* is not due to reduced DNA damage.

DNA-damaging antibiotics are known to induce the bacterial SOS response^36^, a global stress-response regulon, among which *recA* is one of the earliest induced^37,38^. To determine whether deletion of *hsdM* influences SOS induction following ciprofloxacin exposure, we used a P*recA*-*mScarlet* reporter. Using our mother-machine microfluidics platform, we recorded heterogeneous *recA* induction in both the wild type and the Δ*hsdM* strain during exposure to antibiotic-free medium after 4 h exposure to ciprofloxacin at 10× its MIC value. Cells in a non-growing state or undergoing filamentation or lysis displayed low levels of P*recA* fluorescence. In contrast, persisters and cells that doubled before entering in a growth arrested state exhibited high levels of *recA* induction (Fig. 3j, Extended Data Fig. 4h), with the Δ*hsdM* strain displaying a larger proportion of cells with high *recA* induction compared with the PAO1 strain (Fig. 3k). These data suggest that strong SOS activation is associated with survival-related phenotypes during recovery from ciprofloxacin-induced DNA damage and that the loss of *hsdM* increases the probability that individual cells display high SOS activation.

Finally, we introduced a heterologous type II RM system with distinct sequence specificity^25^ into dPAO1 via the pLOLA expression vector^31^ and found that expression of the type II RM system decreased persister levels and lag time, phenocopying complementation with the endogenous type I RM system (Fig. 3l, Extended Data Fig. 4i). Since type I and type II RM systems target different DNA motifs, these data suggest that DNA methylation via a RM system methyltransferase has a profound impact on *P. aeruginosa* PAO1 physiology and its response to DNA-damaging antibiotics and that these effects do not depend on the specific DNA motifs that are methylated.

### Clinical isolates from cystic fibrosis patients lacking type I and type II RM systems display low genetic resistance and high persistence to ciprofloxacin

Next, we asked whether natural variation in RM systems shapes persistence to fluoroquinolones in clinical populations. Understanding these determinants in *P. aeruginosa* is critical given its ecological versatility and the role this pathogen plays in severe infections in hospitalized and immunocompromised patients^39–42^. Clinical isolates frequently encode multiple defence modules, including RM systems^24,43^, suggesting immune architecture may underlie inter-strain variability in persistence to fluoroquinolones.

We analysed 20 clinical isolates from human infections across different infection sites^44,45^(Extended Data Table 4). Genome analysis revealed that six isolates lacked both type I and type II RM systems; interestingly, five of these isolates were from CF patients (Fig. 4a, Extended Data Table 4). Analysis of the complete Bactome collection revealed that only 27% (n = 15) of CF isolates encoded a type I or type II RM system (Extended Data Table 4), consistent with recent evidence^43^. Crucially, these isolates displayed low MIC values (Extended Data Fig. 5a), and high levels of persisters to ciprofloxacin (Fig. 4b and Extended Data Fig. 5b) with an overall negative correlation between persistence and resistance across the 20 clinical isolates (Fig. 4c). Consistent with the data for the PAO1 laboratory strain, increased persister levels were recorded during lag phase but not during exponential growth (Extended Data Fig. 5c). Introducing a functional type I or type II RM system into CF isolates using the pLOLA expression vector^25^ significantly reduced persistence to ciprofloxacin (Fig. 4d). Together, these data suggest that *P. aeruginosa* isolates from the CF lung environment seldomly harbour type I and type II RM systems and that the lack of these systems constrains genetic resistance while increasing persistence to ciprofloxacin.

**Fig. 4.**
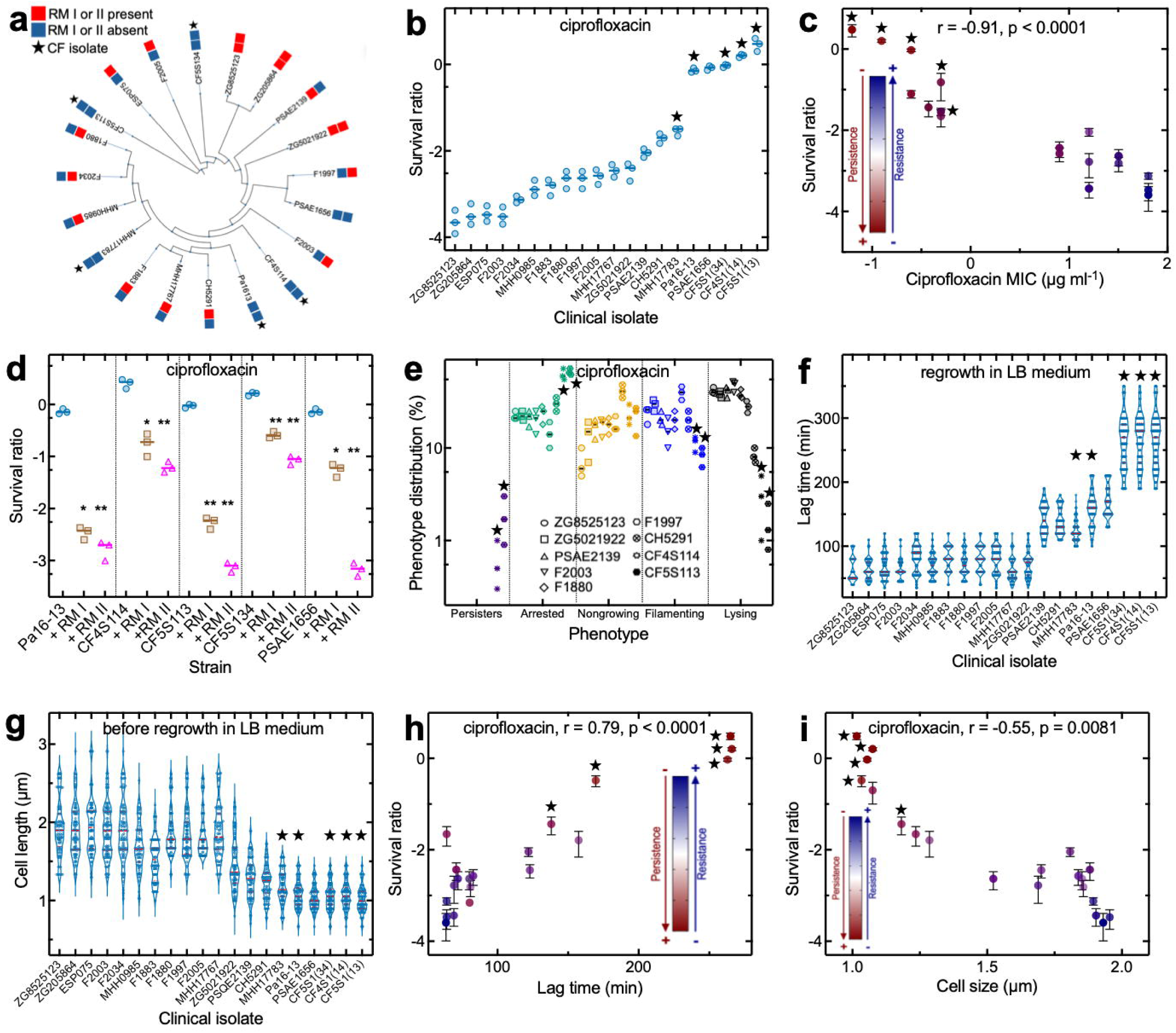
Clinical isolates from cystic fibrosis patients lack RM systems and display slow growth, low genetic resistance and high persistence to ciprofloxacin a, Maximum-likelihood phylogenetic tree for 20 *P. aeruginosa* isolates based on their GyrA protein sequences. Red and blue squares indicate presence or absence of type I (innermost) or II (outermost) RM systems. CF isolates are indicated with stars. The infection source for all the other isolates is reported in Extended Data Table 4. **b,** Corresponding survival ratio of each isolate after exposure to ciprofloxacin at 10× its MIC value and regrowth on plates. Each point represents an independent biological replicate, the centre line indicates the median. **c,** Correlation between the MIC value measured for ciprofloxacin against each isolate and the corresponding survival ratio after exposure to ciprofloxacin at 10× its MIC value. Spearman r = −0.91, p-value < 0.0001. **d,** Survival ratio of the five isolates with the highest persister ratio in the absence and presence of the pLOLA plasmid encoding type I or II RM systems after exposure to ciprofloxacin at 10× its MIC value. Each point represents an independent biological replicate, the centre line indicates the median, statistical significance was determined using Kruskal–Wallis tests with Dunn’s post hoc correction for multi-group analyses; *: p-value: < 0.05; **: p-value: < 0.01. **e,** Quantification of cell-fate distributions for nine clinical isolates spanning the persistence spectrum. Each point represents an independent biological replicate, the centre line indicates the median. **f-g,** Distribution of individual cell lag time and length in the absence of ciprofloxacin for individual cells of each of the 20 clinical isolates. n=150 cells were analysed for each isolate. **h,** Corresponding correlation between the mean lag time and the survival ratio to ciprofloxacin at 10× its MIC value measured for each clinical isolate, Spearman r = 0.79, p-value < 0.0001. **i,** Corresponding correlation between the mean cell length and the survival ratio, Spearman r = −0.55, P = 0.0081). For **c, h** and **i** points are coloured according to persistence and resistance phenotype as indicated by the scale, each point represents the mean of 3 independent replicates and bars denote the corresponding standard deviations.

Single-cell analysis demonstrated that CF isolates exhibited reduced lysis and filamentation, alongside higher levels of persisters and of cells that doubled before entering a growth arrested state compared to isolates from other sites of infection (Fig. 4e). CF isolates also displayed increased lag time and reduced cell length (Fig. 4f, 4g) with an overall positive correlation between lag time and persister levels and a negative correlation between cell length and persister levels (Fig. 4h, 4i), thus reflecting the physiological state induced by RM loss in PAO1. CF isolates also displayed increased doubling time during exponential phase compared to non-CF isolates (Extended Data Fig. 5d). CF isolates did not display increased levels of persisters to the non-DNA-targeting antibiotic ceftazidime and overall lag time did not correlate with levels of persisters to ceftazidime (Extended Data Fig. 5e, 5f), further corroborating that RM-deficiency mediated lag does not confer generic persistence to antibiotics.

## Discussion

The shift that occurs when *P. aeruginosa* moves from the environment into the CF airway involves both nutritional and physicochemical changes for the bacterium, which must survive and adapt to highly stressful conditions^46,47^. Cellular activities and growth rates are drastically changed, with the majority of CF isolates showing a slow growth phenotype which may increase *P. aeruginosa* persistence in the patients^48,49^. However, little is known about the molecular mechanisms leading to this slow growth phenotype in CF isolates^50^. Intriguingly, adaptive laboratory evolution of *P. aeruginosa* CF isolates suggested that reduced growth rate might be linked to altered DNA supercoiling and replication^50^, whereas bioinformatics analysis has shown that *P. aeruginosa* CF isolates are less likely to encode RM systems compared to non-CF patient isolates, possibly due to low phage penetration of the CF airways^28,43^.

Here we introduce a new functional link between the absence of type I and II RM systems and slow growth in *P. aeruginosa* CF isolates, thus reconciling previous observations^28,43,50^ and stimulating new research on the role of RM defence systems in *P. aeruginosa* infections of the CF airways. Considering that *P. aeruginosa* adaptation to the CF airways over hundreds of thousands of generations creates broad phenotypic and genomic changes^46^, we further recapitulated the functional link between the absence of RM systems and slow growth in well-defined mutants of the *P. aeruginosa* PAO1 strain.

The link between slow growth and decreased susceptibility to antibiotics is well established in *E. coli* with persisters to antibiotics within clonal susceptible populations often being in a dormant or slow growing state^30^. Several different processes have been identified to play a role in bacterial persistence to antibiotics, including, stress response signalling molecules^10,11^ DNA damage repair systems and the SOS response^11^, the formation of protein aggregates^51^, reduced antibiotic accumulation^12^, intracellular pH^13^ and toxin-antitoxin systems^6^. Interestingly, toxin-antitoxin systems share some characteristics with RM systems^25^. Comparatively less is known about the impact of the CF slow growth phenotype on the susceptibility of *P. aeruginosa* to antibiotics, which could be crucial when designing treatment of CF infections^50^. Adaptive laboratory evolution of *P. aeruginosa* CF isolates suggested that reduced growth rate decreases susceptibility to bactericidal antibiotics at the population level^50^.

Here we demonstrate that RM-deficient slow growing CF isolates display distinct susceptibility profiles to ciprofloxacin compared to non-CF isolates: lower genetic resistance and higher levels of persistence. It is conceivable that in the stressful conditions encountered in the CF airways^46^, persistence may provide a greater selective advantage than resistance because survival through transient antibiotic exposure allows long-term maintenance of infection reservoirs^6,8,42^. It is also conceivable that the loss of RM-dependent methylation may cause increased DNA supercoiling by gyrase, decreasing the emergence of gyrase mutants resistant to fluoroquinolones in RM-deficient CF isolates. Accordingly, in *Streptococcus pneumoniae* the DpnII and DpnIII RM systems that methylate GATC^4^, i.e. the same DNA sequence methylated by HsdM in *P. aeruginosa*, decrease DNA supercoiling by gyrase and increase the emergence of gyrase mutants resistant to fluoroquinolones^4^.

Moreover, based on our groundbreaking mechanistic data and recent findings in the field of RM systems, we propose the following model to rationalize altered physiology and persistence to fluoroquinolones in RM-deficient *P. aeruginosa* (Extended Data Fig. 6). The loss of RM-dependent methylation causes increased DNA supercoiling by gyrase^4^. Increased DNA supercoiling by gyrase (possibly in concert with nucleoid associated proteins^52^) leads to decreased nucleoid size^53^, chromosomal reorganization and altered gene expression^33,54^ in the absence of RM-dependent methylation. Nucleoid size reduction and increased supercoiling cause a significant delay in DNA replication initiation, as shown for *E. coli*^55^, and in the exit from lag phase compared to RM-proficient *P. aeruginosa*.

Differently from what previously observed for *E. coli*^6,56^, an extended lag phase does not have an impact on persistence to beta-lactams in *P. aeruginosa*. However, slow DNA replication initiation and increased DNA gyrase cleavage^4^, decrease fluoroquinolone-induced collisions between replication forks and gyrase-DNA cleavage complexes in RM-deficient *P. aeruginosa*. Reduced collisions limit lethal chromosome fragmentation and replication fork collapse during exposure to fluoroquinolones^40–44^; however, DNA damage is comparable between RM-deficient or proficient *P. aeruginosa* in accordance with previous reports demonstrating similar DNA damage between cells that survive or are killed by fluoroquinolones^57,58^.

These data are in accordance with recent data obtained for enrofloxacin against *Salmonella enterica*^57^ and are consistent with extensively documented post-antibiotic effects of other antibiotics^59^. Lysis and filamentation is featured also by RM-deficient *P. aeruginosa*; however, these strains also feature larger subpopulations of persisters and cells undergoing doubling before growth arrest. Slow DNA replication initiation and reduced forks-gyrase-DNA collisions allow these cells to activate the SOS response and either survive or delay killing by fluoroquinolones. It is conceivable that cellular heterogeneity in the timing and extent of DNA replication initiation, SOS response to DNA damage and DNA repair could explain heterogeneity in survival to fluoroquinolones in RM-deficient *P. aeruginosa*, as previously proposed for MazF *E. coli* persisters^58^.

In conclusion, our study opens a new area of knowledge in the field of RM biology by demonstrating that loss of RM-dependent methylation profoundly affects *P. aeruginosa* physiology. These findings will stimulate new research in the emergent field of non-canonical cellular functions carried out by defence systems^4,25^. Moreover, the lack of RM systems has a specific impact on *P. aeruginosa* susceptibility to fluoroquinolones that is not further affected by the heterologous introduction of other defence systems. However, it is conceivable that different defence systems play a role in susceptibility to other antibiotics and in different species. In this context, recent seminal work demonstrated that the activation of the CBASS defence system increases *Vibrio cholerae* susceptibility to antifolate antibiotic^2^. Together these findings open a new way of approaching bacterial susceptibility to antibiotics, bringing defence systems in a forward-facing position in this field.

## Methods

### Strains, media, culture conditions and chemicals

Bacterial strains and plasmids used in this study are listed in Extended Data Table 5. All strains are derivatives of *Pseudomonas aeruginosa* repaired MPAO1^60^ (refered as PAO1 in this study or clinical isolates obtained from the Bactome and Weimann collections^44,41^., All experiments were performed at 37 °C in Lysogeny Broth (LB). The following antibiotics: tobramycin, meropenem, ceftazidime, polymyxin B, ofloxacin, moxifloxacin, levofloxacin and ciprofloxacin were prepared from powder (Sigma– Aldrich) and dissolved according to the manufacturer’s instructions.

### Genetic engineering

The dPAO1 strain has been generated as previously described^29^. Briefly, to generate the dPAO1 strain deletions of the candidate defence systems and resident prophages were generated by amplifying upstream and downstream homology arms from the MPAO1 genome. Homology arms ranged from approximately 500 to 2,000 bp depending on the size and structure of the target region. PCR products were assembled into the gentamicin-resistant suicide vector pDONORPex18GmR using NEBuilder HiFi DNA Assembly Master Mix. The resulting plasmids were introduced into *P. aeruginosa* by electroporation using 1 mm gap cuvettes and a Bio-Rad electroporator set to 1.8 kV. Single-crossover integrants were selected on LB agar containing gentamicin at 50 µg ml⁻¹.

Double-crossover mutants were selected on TYS10 agar containing 10% sucrose, as described previously. Candidate sucrose-resistant colonies were screened for gentamicin sensitivity and tested by PCR across the expected deletion junction. PCR products from candidate mutants were confirmed by Sanger sequencing, and final engineered strains were verified by whole-genome sequencing.

To generate deletion of *hsdM* (PA2735) and *hsdR* (PA2732) coding sequences in *P. aeruginosa* PAO1, we used a CRISPR-Cas9 assisted recombineering toolkit^29^. In the case of *hsdR*, considering the overlap upstream with PA2733 coding sequence, a short four codons sequence has been conserved after the start codon of *hsdR*. Deletion of coding sequences have been verified by PCR and sequencing, and curing of the toolkit’s vectors was ensured through loss of resistance (to carbenicillin and streptomycin) and negative PCR.

For complementation of *hsdM*, we amplified its coding sequence and its natural promoter (NOG897/898). For complementation of *hsdR*, we fused the promoter sequence upstream of the operon (NOG899/900) with the coding region of *hsdR* (NOG9001/902) by SOE PCR. The insert was cloned in the pUC18T-mini-Tn7T-Gm plasmid by classic digestion/ligation procedure using SacI/KpnI enzymes. The plasmid was introduced in *P. aeruginosa* together with the pTNS2 plasmid and plated on LB plates supplemented with 30ug/mL Gm. Insertions at the *att*Tn7 site were verified by colony PCR^61^. Finaly, the gentamycin resistance marker was removed through Flp-mediated excision using the pFLP2 plasmid and counter selection on sucrose 5% LB plates. The primers used in this study are listed in Extended Data Table 6.

### Minimum inhibitory concentration (MIC) determination

Minimum inhibitory concentrations (MICs) were determined by broth microdilution as previously described^62^. Briefly, overnight cultures were diluted to a final inoculum of ∼5 × 10⁵ CFU/mL in LB medium. Antibiotics were prepared as twofold serial dilutions in 96-well microtiter plates to cover the appropriate concentration range. Plates were inoculated with bacterial suspensions and incubated at 37 °C for 16 h under aerobic conditions without shaking. MIC values were defined as the lowest antibiotic concentration preventing visible bacterial growth. All measurements were performed with at least three independent biological replicates.

### Time-kill assays

Persistence to antibiotics was assessed by exposing cells to antibiotics at 10× their MIC value for 4 h at 37 °C with shaking as previously described^62^.

All time-kill assays were performed against *P. aeruginosa* in its lag-phase of growth unless otherwise indicated. Overnight cultures (16 h) were diluted 1:100 into LB medium containing antibiotic and immediately incubated in antibiotic in LB medium. Following treatment, cells were pelleted, washed, and resuspended in phosphate-buffered saline (PBS), serially diluted, and plated on LB agar. Colony-forming units (CFUs) were enumerated after 24 h incubation at 37 °C and compared to CFU counts prior to antibiotic exposure. Plates were further incubated for up to 48 h to account for slow-growing survivors.

To confirm persistence, individual survivor colonies were isolated, regrown in antibiotic-free medium, and re-exposed to the same antibiotic conditions as previously described^63^. A persister phenotype was defined by restoration of the original survival fraction and biphasic killing dynamics upon re-treatment^30^.

### Single-cell time-lapse microscopy experiments

Single-cell time-lapse microscopy was performed using our previous described microfluidics-based time-lapse microscopy platform^64^. Specifically, we use a polydimethylsiloxane device^65^ equipped with four identical microfluidic networks that can be controlled simultaneously and independently to maximise experimental throughput. Each of these networks is equipped with approximately 6000 lateral microfluidic channels with width and height of 1 μm each and a length of 20 μm. These lateral channels are connected to a main microfluidic chamber that is 25 μm and 100 μm in height and width, respectively. The device was coated with 5 µl of 0.03% (w/v) chitosan solution for 10 min and subsequently washed with phosphate-buffered saline (PBS). Overnight *P. aeruginosa* cultures were loaded into the device by diffusion, after which the main channel was flushed to remove excess cells as previously described^10^. The microfluidic chip was connected to fluorinated ethylene propylene tubing and mounted on an inverted microscope housed within an environmental chamber maintained at 37 °C, as described previously^66^. LB medium with or without antibiotics was continuously infused at a flow rate of 60 mbar. Images were acquired every 5 min for a total duration of 8 h for each experiment^67^.

Cell lineages were reconstructed by tracking cells over time and manually annotating division events. Lag time for each cell was defined as the time from loading into the mother machine after overnight culture to the first division event of the cell. Division time was defined as the interval between cell birth and subsequent division.

For P*recA-mscarlet*, fluorescence was measured at each time point using TRITC filter (excitation 550nm, emission 580nm) with an exposure of 0.06 ms.

### Phage infection

To measure the effect of RM type I on phage infection, the dPAO1 strain was transformed with the pLoLa plasmid encoding the RM type I system under its native promoter, or mCherry gene for control. Cultures were grown in LB medium to logarithmic phase, diluted to an OD_600_ of 0.03, and mixed with phage at a multiplicity of infection (MOI) of 0.1. Bacterial growth was monitored by measuring the OD_600_ over 15 h at 37 °C in a plate reader.

Growth in the presence or absence of phage was quantified from the OD_600_ time-course data. For each strain-phage combination, the area under the growth curve (AUC) was calculated between 3 and 10 hours post infection, and these values were used as an estimate of bacterial growth. For heatmap visualisation, AUC values from all experimental runs were normalised to the mean AUC of the mCherry no phage control.

Statistical analyses were performed separately for each phage using least-squares linear models, with normalised AUC as the response variable and strain as a categorical predictor. mCherry was used as the reference strain. p-values for strain effects were corrected for multiple testing across all phage-strain comparisons using the Benjamini-Hochberg false discovery rate procedure. Asterisks in the heatmap indicate strains with significantly higher growth than mCherry after correction (adjusted p < 0.05).

### Transcriptomic analysis

RNA extraction for RNAseq was performed using the RNeasy Mini Kit (Qiagen) in combination with Qiashredder™ columns as described previously^68^. Briefly, 10 ml bacterial cultures were inoculated with a starting OD_600_ of 0.05, harvested after 17 h and mixed with an equal volume of RNA protect (Qiagen). The same cultures were diluted at a 1:10 ratio (v/v) in preheated LB medium and grown for 1 h. After RNA extraction, DNA removal was performed using the DNA-free™ Kit (Thermo Fisher Scientific, Waltham, MA, USA). RNA integrity was checked using Fragment Analyzer (5300 Fragment analyzer system; Agilent). Ribosomal RNA was removed using the *Pseudomonas aeruginosa* Ribopool kit (siTOOLS Biotech) and sequencing of the samples was performed in paired-end mode (2 × 50 bp reads) on an Illumina NovaSeq 6000 device. Sequencing reads were mapped to the PAO1 reference genome using bowtie2^69^, and the number of reads per gene was assessed with FeatureCounts^70^. Differential gene expression analysis between the PAO1, Δ*hsdR* and Δ*hsdM* strains was performed with the edgeR package^71^ using the function glmTreat (fold-change 1.2). Genes were filtered using the edgeR function filterByExpr, and reads were normalized using the edgeR function calcNormFactors (trimmed mean of *M* values). Adjusted *P*-values were calculated for multiple tests using the Benjamini–Hochberg adjustment. The significance threshold was set to a false discovery rate (FDR) of ≤ 0.05 to identify differentially expressed genes. Functional enrichment analysis of differentially expressed genes was performed using DAVID (Database for Annotation, Visualization and Integrated Discovery). Gene lists were analysed against Gene Ontology (GO) terms and KEGG pathways, using the full genome as background. Enriched terms were identified based on modified Fisher’s exact test (*EASE score*) and corrected for multiple testing using the Benjamini–Hochberg procedure. Terms with an adjusted *P* value < 0.05 were considered significantly enriched.

### Chromosome equivalent analysis

Chromosome equivalent quantification was performed as previously described^34^. Briefly, overnight cultures were diluted 1:1000 in LB medium and incubated at 37 °C with agitation until OD_600_ reached 0.05. Aztreonam (10 μg ml^−1^) and rifampicin (200 μg ml^−1^) were added simultaneously to respectively block cell division and inhibit new initiation of replication cycle. Rifampicin inhibits initiation of replication but allows ongoing replication rounds to finish, thus enabling estimation of the number of chromosome equivalents. Cells were incubated simultaneously with both antibiotics at 37 °C for 120 min. Cells were washed with PBS before being fixed with 100% cold (4°C) methanol for 1 hour. Fixed cells were washed with PBS, and the DNA was stained with DAPI (4′,6-diamidino-2-phenylindole; 0.3 μg ml^−1^). For each time point, a minimum of 50,000 cells was analyzed using a Cytoflex flow cytometer with a maximum of 2000 events counted per second as previously described^12^. DAPI was excited with an ultraviolet laser at 355 nm and emission recorded at 450 nm. The number of the peaks in each histograms show the chromosome equivalents per cell, i.e. the number of fully replicated chromosomes and reflects the number of origins present in the cell at the time the drugs were added. Number of genome equivalents (*N*) was calculated using stationary phase cells (1*N*) as a reference.

### Transmission Electron Microscopy

For transmission electron microscopy (TEM) analysis, cells were fixed in 2% glutaraldehyde and 2% paraformaldehyde in 0.1 M piperazine (PIPES) buffer, pH 7.2 for 1 h at room temperature, then washed 3 times for 5 min with PIPES buffer. Samples were post-fixed in 1% aqueous osmium tetroxide, reduced in 1.5% potassium ferrocyanide for 1 h, then washed 3 times for 5 min in deionised water before dehydration in a graded ethanol series (30%, 50%, 70%, 80%, 90%, 95% for 10 min each, then 2 times for 15 min in 100% ethanol). Cells were gradually embedded in Durcupan resin (Sigma Aldrich, Merck, Gillingham, UK) over 24 h, then polymerised in BEEM capsules at 62 degrees for 24 h. 70 nm ultrathin sections were collected on formvar-coated copper mesh grids, contrasted in lead citrate and were imaged using a JEOL JEM 1400 transmission electron microscope operated at 120 kV equipped with a digital camera (Gatan Rio16, Ametek, Leicester, UK).

### Nucleoid fraction quantification

Nucleoid occupancy was quantified from TEM images based on grayscale intensity profiles. Individual cells were manually segmented, and a line profile was drawn along the longitudinal axis of each cell, from pole to pole. Grayscale intensity profiles were extracted and smoothed using a Gaussian blur (σ = 2 pixels) to reduce high-frequency noise.

The nucleoid was identified as a relatively electron-lucent region with higher grayscale intensity than the surrounding, ribosome-rich cytoplasm. The cytoplasmic baseline intensity (μ_cyto) was defined as the mean grayscale intensity measured at the two cell poles, where nucleoid material was largely absent. The standard deviation of the cytoplasmic intensity (σ_cyto) was calculated from these regions. A threshold intensity (T) was defined as T = μ_cyto + 2σ_cyto. Regions with grayscale intensities above this threshold were classified as nucleoid-associated regions.

Nucleoid occupancy was calculated as the proportion of the longitudinal cell profile exceeding this threshold relative to the total cell length. This analysis provided a measure of the fraction of the cell occupied by a distinguishable nucleoid-associated region.

### Chromosome condensation index

Local DNA condensation was quantified from TEM images using Fourier transform– based spatial frequency analysis. Individual cells were manually segmented, and nucleoid regions were selected for analysis.

Grayscale images were normalized and denoised prior to analysis. Two-dimensional fast Fourier transforms (FFT) were computed for each nucleoid region using ImageJ or equivalent software. The resulting power spectrum *P*(*r*) was radially averaged, where *r* represents the radial distance from the center of the Fourier space. The maximum radius *R*_max_ corresponds to the highest spatial frequency captured in the image (Nyquist limit).

A normalized condensation index (CI) was calculated as the ratio of high-frequency to low-frequency spectral power:

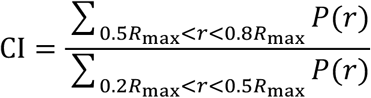

We separated the Fourier spectrum into low-(0.2–0.5) and high-frequency (0.5–0.8) components to distinguish large-scale chromosome organization from local finer-scale structural features^72^. The use of normalized radial frequency ranges enables comparison across images independent of resolution and sampling.

Condensation indices were calculated for multiple cells per condition and averaged across biological replicates. Comparable CI values indicate similar local DNA packing despite differences in global chromosome organization.

### Quantification of DNA damage and cell integrity

DNA damage was quantified using the Promega DeadEnd™ Fluorometric TUNEL system, which labels DNA strand breaks by terminal deoxynucleotidyl transferase (TdT)-mediated incorporation of fluorescein-dUTP at free 3′-OH ends^73^.

Overnight cultures (16 h) were diluted 1:100 into LB medium containing ciprofloxacin at either 1× or 10× its MIC value and incubated for 4 h at 37 °C with shaking. Cells were harvested by centrifugation (5,000 rpm, 10 min, 4 °C), washed twice with cold phosphate-buffered saline (PBS), and resuspended in 0.5 ml PBS.

Cells were fixed in 1% methanol-free formaldehyde on ice for 60 min, washed twice with cold PBS, and permeabilized for 20 min according to the manufacturer’s protocol. Following two additional PBS washes, TUNEL staining was performed according to the manufacturer’s instructions.

Fluorescence was measured using a CytoFLEX flow cytometer as previously described^12^, and at least 50,000 cells were analysed per sample, with acquisition rates not exceeding 2,000 events per second.

Cell integrity was evaluated by forward scatter (FSC) and side scatter (SSC) parameters, which reflect cell size and granularity, respectively. A gate was established based on the FSC–SSC profile of the non-treated overnight culture to exclude debris and dead cells. Only events falling within this gate were included in the intact cells count.

### Phylogenetic analysis

The GyrA (DNA gyrase A subunit) protein sequences from clinical isolates were used to infer phylogenetic relationships. Amino acid sequences were aligned using MEGA X with the MUSCLE algorithm. The best-fit substitution model was determined in MEGA, and a Maximum Likelihood (ML) tree was constructed with 1,000 bootstrap replicates to assess branch support. Trees were generated as unrooted and exported in Newick format. Final tree visualization and annotation were performed using iTOL (https://itol.embl.de).

### Detection of defence systems in clinical isolates

Genomes of clinical isolates were screened for bacterial defence systems using PADLOC^74^ and **DefenseFinder**^75^. Genomic assemblies in FASTA format were analysed with default parameters to identify known antiviral and other defence systems, including RM systems, CRISPR-Cas, BREX, DISARM, and recently described systems. Predicted defense elements were curated to retain only complete operons or essential signature genes.

### Statistical analysis

All analyses were done using GraphPad Prism 10 (GraphPad Software, Inc., La Jolla, CA).

## Data availability

All sequencing data analysed in this study are publicly available. The raw sequence reads are deposited in the **NCBI Sequence Read Archive (SRA)** under BioProject accession **PRJNA526797** and can be accessed via the NCBI SRA database. Any other datasets generated during the study (e.g. processed datasets, analysis outputs, or supplementary source files) are included within the manuscript and supplementary information files. Further materials supporting the findings of this study are available from the corresponding author upon reasonable request.

The raw transmission electron microscopy (TEM) images and supplementary movie files supporting the findings of this study have been deposited in the Figshare repository under the collection DOI: https://figshare.com/s/5bf31929186cbef619f2. These datasets are publicly available as of the date of publication.

## Funding

This work was supported by the BBSRC and the EPSRC through two grants awarded to S.P. and U.L. (BB/V008021/1, EP/Y023528/1). This work was further supported via the JPIAMR project ERADIAMR (MR/Y033892/1) awarded to S.P., R.C. and S.H. and the BBSRC project MULTI-DEFENCE (BB/X003051/1) awarded to E.R.W., S.v.H. and S.P. S.H. was also funded by the Novo Nordisk Foundation (NNF 18OC0033946), and received funding from the Deutsche Forschungsgemeinschaft (DFG, German Research Foundation) under Germany’s Excellence Strategy – EXC 2155 “RESIST” – Project ID 390874280, within the SFB/TRR-298-SIIRI – Project-ID 426335750 and in the SPP 2389 (HA 3299/9-1, AOBJ: 687646). The funders had no role in study design, data collection and analysis, decision to publish, or preparation of the manuscript. The views expressed are those of the authors. For the purpose of open access, the authors have applied a ‘Creative Commons Attribution (CC BY) licence to any Author Accepted Manuscript version arising from this submission.

## Author contributions

Conceptualization, M.M. and S.P.; methodology, M.M., N.O.G., A.O., A.A., U.L., M.P., M.G-C., C.H., R.C., S.v.H., S.H., E.R.W., and S.P.; formal analysis, M.M., N.O.G., A.O., A.A., U.L., M.P., M.G-C., C.H., R.C., S.v.H., S.H., E.R.W., and S.P.; generation of figures, M.M., N.O.G. and S.P.; investigation, M.M., N.O.G., A.A., C.H.; resources, M.M., N.O.G., A.O., A.A., U.L., M.P., M.G-C., C.H., R.C., S.v.H., S.H., E.R.W., and S.P.; data curation, M.M., N.O.G., A.A. and S.P.; writing – original draft, M.M. and S.P.; writing – review & editing, M.M., N.O.G., A.O., A.A., U.L., M.P., M.G-C., C.H., R.C.,S.v.H., S.H., E.R.W., and S.P.; visualization, M.M., N.O.G., and S.P.; supervision, R.C., S.v.H., S.H., E.R.W., and S.P.; project administration, S.P.; funding acquisition, U.L., R.C., S.v.H., S.H., E.R.W., and S.P..

## Competing financial interests

The authors declare no competing financial interests.

## Extended Data legends

**Extended Data Table 1** Length of individual cells for the PAO1, dPAO1, Δ*hsdMSR*, Δ*hsdM*, Δ*hsdM::hsdM*, Δ*hsdR* and Δ*hsdR::hsdR* strains.

**Extended Data Table 2** Lag time of individual cells for the PAO1, dPAO1, Δ*hsdMSR*, Δ*hsdM*, Δ*hsdM::hsdM*, Δ*hsdR* and Δ*hsdR::hsdR* strains.

**Extended Data Table 3** Generation time of individual cells for the PAO1, dPAO1, Δ*hsdMSR*, Δ*hsdM*, Δ*hsdM::hsdM*, Δ*hsdR* and Δ*hsdR::hsdR* strains.

**Extended Data Table 4** Clinical isolates name, site of infection, and presence of type I or type II RM systems.

**Extended Data Table 5** Strains and Plasmids used in this study.

**Extended Data Table 6** Primers used in this study.

**Extended Data Fig. 1 | Deletion of the type I RM system increases persistence to fluoroquinolones without genetic resistance.**

**a,** Minimum inhibitory concentration (MIC) values measured for tobramycin, meropenem, ceftazidime, polymyxin B, ciprofloxacin, ofloxacin, moxifloxacin and levofloxacin against the PAO1 and dPAO1 strains indicated in black and blue, respectively. Each point represents an independent biological replicate, the centre line indicates the median.

**b-e,** Temporal dependence of the survival ratio of the PAO1 and dPAO1 strains during exposure to **b,** tobramycin, **c,** ceftazidime, **d,** polymyxin B and **e,** ciprofloxacin at 10× their respective MIC value. Black and blue circles represent data for the first treatment against the PAO1 and dPAO1 strains, respectively, whereas the yellow and green circles represent the corresponding data for the second treatment. Each circle and error bar represents the mean and standard deviation of values obtained in biological replicates. Lines are guide-for-the-eyes.

**f,** Survival ratio of the PAO1 and dPAO1 strains after 4 h exposure to ciprofloxacin, ofloxacin, moxifloxacin or levofloxacin and regrowth on plates. Each point represents an independent biological replicate, the centre line indicates the median. Statistical significance was determined using two-tailed Mann–Whitney U tests vs the PAO1 strain; ****: p-value < 0.0001.

**Extended Data Fig. 2 | Phage resistance conferred by the type I RM system.**

**a, b,** Heat maps showing the relative growth of dPAO1 strains complemented either with the pLOLA plasmid encoding type I RM system (RMI, left) or with the empty pLOLA-mCherry control plasmid (right) following infection with **(a)** lytic or **(b)** temperate bacteriophages. The area under the growth curve (AUC) was calculated between 3 and 10 hours post infection, and these values were used as an estimate for bacterial growth. AUC values from all experimental runs were normalised to the mean AUC of the mCherry no phage control. Statistical analyses were performed separately for each phage using least-squares linear models, with normalised AUC as the response variable and strain as a categorical predictor. mCherry was used as the reference strain. p-values for strain effects were corrected for multiple testing across all phage-strain comparisons using the Benjamini-Hochberg false discovery rate procedure. Color intensity represents the relative bacterial growth after phage exposure, with blue indicating resistance and red indicating susceptibility. Asterisks in the heatmap indicate that the type I RM complemented strain displayed significantly higher growth than mCherry strain after correction, i.e. adjusted p-value < 0.05.

**Extended Data Fig. 3 | Working principle of the type I RM system.**

Cartoon illustrating the role of the type I RM system in DNA methylation and cleavage. The methylation complex (HsdM–HsdS) modifies host DNA, enabling recruitment of HsdR to form the active RM complex. In the absence of appropriate methylation, unmethylated DNA is targeted for ATP-dependent cleavage.

**Extended Data Fig. 4 | Type I RM activity influences the nucleoid dimension, chromosome replication and SOS response.**

**a,** Volcano plot of differential gene expression between the Δ*hsdM* and the PAO1 parental strain during stationary phase, after 17 h of growth in LB medium. Significantly up-or down-regulated genes are indicated in red and blue, respectively.

**b**, Heatmap showing the log₂ fold change (log₂FC) in the expression of each significantly differentially regulated gene (locus tags indicated) in the Δ*hsdM* strain relative to the PAO1 strain during lag phase. The colour intensity reports the magnitude of log₂FC, with red and blue indicating upregulation and downregulation, respectively.

**c,** DAPI fluorescence intensity in individual cells of the PAO1 strain in stationary phase and the Δ*hsdR* and Δ*hsdMSR* strains in lag phase after 120 min of incubation with rifampicin (200μg/mL) and aztreonam (10μg/mL). The results are representative of three independent experiments each measuring N=50,000 individual cells. 1N and 2N indicate 1 and 2 number of chromosome equivalents, respectively.

**d,** Representative fast Fourier transform (FFT) analysis of grey intensity signal profiles obtained from electron microscopy images showing normalized integrated density as a function of the nucleoid radius for the PAO1, Δ*hsdR*, Δ*hsdM* and Δ*hsdMSR* strains. **e,** Representative forward scatter (FSC-H) versus side scatter (SSC-H) density plots for the PAO1 and Δ*hsdM* strains treated with ciprofloxacin at 10× its MIC value. The polygon gate identifies the intact bacterial cells based on measurements on these same strains but without treatment with ciprofloxacin.

**f,** Distribution of TUNEL staining fluorescence for the PAO1, Δ*hsdM*, Δ*hsdR*, Δ*hsdM* and Δ*hsdMSR* strains after exposure to ciprofloxacin at 1× (left) and 10× (right) its MIC value. Brackets denote the fluorescence gate used to quantify the FITC-positive population.

**g,** Corresponding percentage of cells displaying DNA strand breaks after exposure to ciprofloxacin at 1× its MIC value. Each point represents an independent biological replicate measuring N=50,000 individual cells, the centre line indicates the median. **h,** Temporal dynamics of P*recA-mscarlett* fluorescence during and after treatment with ciprofloxacin at 10× its MIC value stratified by cell fate for the PAO1 strain. Points and shaded areas represent the mean and standard deviation of measurements carried out for N=30 individual cells per phenotype.

**i,** Distribution of lag times of individual cells for the PAO1, dPAO1 and dPAO1 strain complemented with the pLOLA plasmid encoding type I or II RM systems. The red lines indicate the median and quartiles. Statistical significance was determined using two-tailed Mann–Whitney U tests vs the PAO1 strain, ****: p-value < 0.0001. N=229, 139, 132, 205 for the PAO1, dPAO1, dPAO1 + type I RM and dPAO1 + type II RM strains.

**Extended Data Fig. 5 | The absence of type I and II RM systems drives slow growth and higher persistence to ciprofloxacin in CF clinical isolates of *P. aeruginosa*.**

**a,** MIC values measured for ciprofloxacin against 20 *P. aeruginosa* clinical isolates in biological triplicate, stars indicate CF isolates.

**b,** Temporal dependence of the survival ratio of the PSA1656 and ZG8525123 isolates during exposure to ciprofloxacin at 10× its MIC value. Blue triangles and black circles represent data for the first treatment against the PSA1656 and ZG8525123 isolates, respectively, whereas the yellow and green points represent the corresponding data for the second treatment. Each circle and bar represents the mean and standard deviation of values obtained in biological replicates. Lines are guides-for-the-eye. **c,** Temporal dynamics of survival ratio of the CF5S1(13), CF5S1(14), PSAE1656, ZG8525123 and F1880 cultures exposed at the indicated time points to ciprofloxacin at 10× its MIC value and regrowth on antibiotic-free plates. Points represent the mean and standard deviation of three independent biological replicates.

**d,** Distribution of doubling times of individual cells for 20 clinical isolates. Red lines indicate the median and quartiles of each distribution with values measured for N=150 cells for each isolate.

**e,** Survival ratio of each isolate after exposure to ceftazidime at 10× its MIC value and regrowth on plates. Each point represents an independent biological replicate, the centre line indicates the median.

**f**, Corresponding correlation between the mean lag time and the survival ratio after exposure to ceftazidime at 10× its MIC value for 4 h measured for each clinical isolate. Points are coloured according to persistence and resistance phenotype as indicated by the scale bar. Each point represents the mean and standard deviation of 3 independent replicates, Spearman correlation analysis shows a non-significant association with r = 0.30.

**Extended Data Fig. 6 | Proposed working principle for methylation-dependent, chromosome replication dynamics and persistence to fluoroquinolones.**

In HsdM-proficient *P. aeruginosa*, active replication initiation and replication during regrowth increase the likelihood of replication forks colliding with ciprofloxacin-poisoned gyrase complexes. These collisions promote replication fork collapse, double-strand DNA breaks, chromosome disintegration and cell death, resulting in populations enriched in filamenting and lysed cells following exposure to DNA-targeting antibiotics. In contrast, HsdM-deficient *P. aeruginosa* exhibits altered nucleoid organization together with reduced replication initiation and slower replication dynamics during lag phase and recovery. Under these conditions, the SOS response is activated after ciprofloxacin removal further limiting catastrophic replication-associated chromosome fragmentation. This replication-restrained physiological state promotes growth arrest and survival, leading to enrichment of persister cells. Although both populations accumulate DNA damage, RM-deficient *P. aeruginosa* retains a larger proportion of structurally intact cells, suggesting that chromosome organization and replication dynamics, rather than DNA break formation itself, determine survival outcomes following fluoroquinolone treatment.

## Notes

### Competing Interest Statement

The authors have declared no competing interest.

