## Extended Table 1 for "Loss of restriction-modification methyltransferases drives persistence to fluoroquinolones in *Pseudomonas aeruginosa*"

|  | **PAO1** | **dPAO1** | **PAO1 Δ*hsdMSR*** | **PAO1 Δ*hsdM*** | **PAO1 Δ*hsdR*** | **PAO1 Δ*hsdM::hsdM*** | **PAO1 Δ*hsdR::hsdR*** |
| --- | --- | --- | --- | --- | --- | --- | --- |
| Number of cells | 231 | 224 | 225 | 246 | 231 | 247 | 224 |
| Minimum | 0.8662 | 0.5983 | 0.6286 | 0.5983 | 0.8462 | 1.129 | 1.198 |
| 25% Percentile | 1.321 | 1.024 | 1.100 | 1.160 | 1.301 | 1.611 | 1.648 |
| Median | 1.568 | 1.189 | 1.242 | 1.338 | 1.548 | 1.757 | 1.802 |
| 75% Percentile | 1.812 | 1.301 | 1.392 | 1.511 | 1.792 | 1.923 | 1.952 |
| Maximum | 2.505 | 1.757 | 1.892 | 2.122 | 2.485 | 2.392 | 2.492 |
| Range | 1.638 | 1.158 | 1.264 | 1.524 | 1.638 | 1.264 | 1.294 |
| 5% Percentile | 1.068 | 0.8012 | 0.8678 | 0.8538 | 1.048 | 1.379 | 1.468 |
| 95% Percentile | 2.209 | 1.482 | 1.694 | 1.880 | 2.189 | 2.231 | 2.186 |
| Mean | 1.594 | 1.162 | 1.248 | 1.332 | 1.574 | 1.772 | 1.813 |
| Std. Deviation | 0.3471 | 0.2080 | 0.2344 | 0.2824 | 0.3471 | 0.2445 | 0.2305 |
| Std. Error of Mean | 0.02284 | 0.01390 | 0.01563 | 0.01801 | 0.02284 | 0.01556 | 0.01540 |
| Coefficient of variation | 21.78% | 17.89% | 18.79% | 21.19% | 22.06% | 13.80% | 12.72% |

**Extended Data Table 1** Length of individual cells for the PAO1, dPAO1, Δ*hsdMSR*, Δ*hsdM*, Δ*hsdM::hsdM*, Δ*hsdR* and Δ*hsdR::hsdR* strains.
