## Extended Table 2 for "Loss of restriction-modification methyltransferases drives persistence to fluoroquinolones in *Pseudomonas aeruginosa*"

|  | **PAO1** | **dPAO1** | **PAO1 Δ*hsdMSR*** | **PAO1 Δ*hsdM*** | **PAO1 Δ*hsdR*** | **PAO1 Δ*hsdM::hsdM*** | **PAO1 Δ*hsdR::hsdR*** |
| --- | --- | --- | --- | --- | --- | --- | --- |
| **Number of cells** | 225 | 227 | 226 | 226 | 220 | 223 | 223 |
| **Minimum** | 45.00 | 90.00 | 90.00 | 90.00 | 60.00 | 45.00 | 45.00 |
| **25% Percentile** | 60.00 | 120.0 | 120.0 | 120.0 | 90.00 | 60.00 | 60.00 |
| **Median** | 60.00 | 130.0 | 130.0 | 130.0 | 100.0 | 60.00 | 60.00 |
| **75% Percentile** | 75.00 | 140.0 | 140.0 | 140.0 | 110.0 | 80.00 | 75.00 |
| **Maximum** | 110.0 | 180.0 | 180.0 | 180.0 | 150.0 | 130.0 | 110.0 |
| **Range** | 65.00 | 90.00 | 90.00 | 90.00 | 90.00 | 85.00 | 65.00 |
| **5% Percentile** | 45.00 | 100.0 | 100.0 | 100.0 | 80.00 | 60.00 | 45.00 |
| **95% Percentile** | 80.00 | 160.0 | 160.0 | 160.0 | 130.0 | 90.00 | 90.00 |
| **Mean** | 62.87 | 130.4 | 131.1 | 130.3 | 101.1 | 69.98 | 65.83 |
| **Std. Deviation** | 13.92 | 16.56 | 17.64 | 17.19 | 18.32 | 13.89 | 12.83 |
| **Std. Error of Mean** | 0.9277 | 1.099 | 1.173 | 1.143 | 1.235 | 0.9301 | 0.8595 |
| **Coefficient of variation** | 22.14% | 12.70% | 13.46% | 13.19% | 18.13% | 19.85% | 19.50% |

**Extended Data Table 2** Lag time of individual cells for the PAO1, dPAO1, Δ*hsdMSR*, Δ*hsdM*, Δ*hsdM::hsdM*, Δ*hsdR* and Δ*hsdR::hsdR* strains.
