## Extended Table 3 for "Loss of restriction-modification methyltransferases drives persistence to fluoroquinolones in *Pseudomonas aeruginosa*"

|  | **PAO1** | **dPAO1** | **PAO1 Δ*hsdMSR*** | **PAO1 Δ*hsdM*** | **PAO1 Δ*hsdR*** | **PAO1 Δ*hsdM::hsdM*** | **PAO1 Δ*hsdR::hsdR*** |
| --- | --- | --- | --- | --- | --- | --- | --- |
| **Number of cells** | 229 | 231 | 237 | 240 | 228 | 229 | 277 |
| **Minimum** | 20.00 | 15.00 | 15.00 | 15.00 | 15.00 | 20.00 | 15.00 |
| **25% Percentile** | 30.00 | 30.00 | 30.00 | 30.00 | 30.00 | 30.00 | 30.00 |
| **Median** | 35.00 | 35.00 | 35.00 | 35.00 | 35.00 | 35.00 | 35.00 |
| **75% Percentile** | 40.00 | 40.00 | 40.00 | 40.00 | 40.00 | 40.00 | 40.00 |
| **Maximum** | 55.00 | 80.00 | 65.00 | 60.00 | 80.00 | 80.00 | 80.00 |
| **Range** | 35.00 | 65.00 | 50.00 | 45.00 | 65.00 | 60.00 | 65.00 |
| **5% Percentile** | 20.00 | 20.00 | 20.00 | 20.00 | 20.00 | 20.00 | 20.00 |
| **95% Percentile** | 50.00 | 60.00 | 50.50 | 50.00 | 50.00 | 50.00 | 60.00 |
| **Mean** | 34.87 | 36.75 | 35.13 | 35.81 | 34.56 | 35.00 | 37.65 |
| **Std. Deviation** | 7.418 | 12.07 | 9.978 | 8.967 | 10.62 | 8.030 | 11.70 |
| **Std. Error of Mean** | 0.4902 | 0.7942 | 0.6481 | 0.5788 | 0.7034 | 0.5306 | 0.7029 |
| **Coefficient of variation** | 21.27% | 32.84% | 28.41% | 25.04% | 30.73% | 22.94% | 31.07% |

**Extended Data Table 3** Generation time of individual cells for the PAO1, dPAO1, Δ*hsdMSR*, Δ*hsdM*, Δ*hsdM::hsdM*, Δ*hsdR* and Δ*hsdR::hsdR* strains.
