## Extended Table 4 for "Loss of restriction-modification methyltransferases drives persistence to fluoroquinolones in *Pseudomonas aeruginosa*"

| **Isolate name** | **Site of infection** | **RM type I** | **RM type II** | **CF isolate** |
| --- | --- | --- | --- | --- |
| PSAE1656 | Respiratory tract | No | No | No |
| F2034 | Rectal smear | Yes | No | No |
| ZG5021922 | Respiratory tract | Yes | Yes | No |
| ZG205864 | Urinary tract | Yes | Yes | No |
| ZG8525123 | Urinary tract | Yes | Yes | No |
| F1880 | Respiratory tract | Yes | No | No |
| F1883 | Respiratory tract | Yes | No | No |
| F2003 | Urinary tract | No | Yes | No |
| MHH0985 | Other | Yes | No | No |
| MHH17767 | Respiratory tract | Yes | No | No |
| F1997 | Rectal smear | No | Yes | No |
| CH5291 | Rectal smear | Yes | No | No |
| PSAE2139 | Urinary tract | Yes | No | No |
| F2005 | Urinary tract | No | Yes | No |
| ESP075 | Other | No | Yes | No |
| Pa16-13 | Respiratory tract | No | No | Yes |
| MHH17783 | Respiratory tract | No | No | Yes |
| CF4S1(14) | Respiratory tract | No | No | Yes |
| CF5S1(13) | Respiratory tract | No | No | Yes |
| CF5S1(34) | Respiratory tract | No | No | Yes |
| CF592_Iso2 | Respiratory tract | No | No | Yes |
| MHH16530 | Respiratory tract | No | Yes | Yes |
| MHH16563 | Respiratory tract | No | Yes | Yes |
| CF609_Iso3 | Respiratory tract | Yes | No | Yes |
| MHH15204 | Respiratory tract | No | No | Yes |
| ZG5003493 | Respiratory tract | No | No | Yes |
| MHH16951 | Respiratory tract | No | No | Yes |
| ZG5089456 | Respiratory tract | Yes | Yes | Yes |
| MHH17546 | Respiratory tract | Yes | No | Yes |
| CH2678 | Respiratory tract | No | No | Yes |

**Extended Data Table 4** Clinical isolates name, site of infection, and presence of type I or type II RM systems.
