## Extended Table 5 for "Loss of restriction-modification methyltransferases drives persistence to fluoroquinolones in *Pseudomonas aeruginosa*"

| **Strain** | **Description** | **Reference** |
| --- | --- | --- |
| ***Pseudomonas aeruginosa*** | | |
| PAO1 WT | Wild-Type strain | 1 |
| dPAO1 | PAO1 deletion of all three chromosomally encoded defence systems—Helicase–DUF2290, Gabija and a type I restriction–modification (RM) system—as well as the prophages Pf1 and phiCTX | ^29^ |
| dPAO1/pLOLA-Rst Helicase DUF2290 | dPAO1 carying a pLOLA plasmid encoding for the Rst Helicase DUF2290 defence system present in PAO1 | This study |
| dPAO1/pLOLA-Gabija | dPAO1 carying a pLOLA plasmid encoding for the Gabija defence system present in PAO1 | This study |
| dPAO1*/*pLOLA-RM type I | dPAO1 carying a pLOLA plasmid encoding for the RM type I defence system present in PAO1 | This study |
| dPAO1*/*pLOLA-RM type II | dPAO1 carying a pLOLA plasmid encoding for the RM type II defence system present in PAO1 | This study |
| dPAO1/pLOLA-MANTIS | dPAO1 carying a pLOLA plasmid encoding a PAMS defence system | This study |
| dPAO1/pLOLA-Pycsar | dPAO1 carying a pLOLA plasmid encoding a Pycsar defence system | This study |
| dPAO1/pLOLA-McPol | dPAO1 carying a pLOLA plasmid encoding a McPol defence system | This study |
| dPAO1/pLOLA-CBASS type I | dPAO1 carying a pLOLA plasmid encoding a CBASS type I defence system | This study |
| dPAO1/pLOLA-CBASS type II | dPAO1 carying a pLOLA plasmid encoding a CBASS type II defence system | This study |
| dPAO1/pLOLA-Shango | dPAO1 carying a pLOLA plasmid encoding a Shango defence system | This study |
| dPAO1/pLOLA-Lamassu | dPAO1 carying a pLOLA plasmid encoding a Lamassu defence system | This study |
| dPAO1/pLOLA-Avast | dPAO1 carying a pLOLA plasmid encoding a Avast defence system | This study |
| dPAO1/pLOLA-Kiwa | dPAO1 carying a pLOLA plasmid encoding a Kiwa defence system | This study |
| dPAO1/pLOLA-Septu | dPAO1 carying a pLOLA plasmid encoding a Septu defence system | This study |
| dPAO1/pLOLA-SEFIR | dPAO1 carying a pLOLA plasmid encoding a SEFIR defence system | This study |
| dPAO1/pLOLA-qatABCD | dPAO1 carrying a pLOLA plasmid encoding a qatABCD defence system | This study |
| PAO1 Δ4 *hsdMSR+* | PAO1 deletion of two chromosomally encoded defence systems—Helicase–DUF2290, Gabija—as well as the prophages Pf1 and phiCTX | This study |
| PAO1 Δ*hsdMSR* | PAO1 deleted for the entire *hsdMSR* operon | ^33^ |
| PAO1 Δ*hsdM* | PAO1 CRISPR-Cas9 deletion of *PA2735* (*hsdM*) | This study |
| PAO1 Δ*hsdM*::*hsdM* | PAO1 CRISPR-Cas9 deletion of *PA2735* (*hsdM*); chromosomal insertion at *att*Tn7 site of *hsdM* sequence under control of its natural promoter (NOG897/898) | This study |
| PAO1 Δ*hsdR* | PAO1 CRISPR-Cas9 deletion of *PA2732* (*hsdR*) | This study |
| PAO1 Δ*hsdR*::*hsdR* | PAO1 CRISPR-Cas9 deletion of *PA2732* (*hsdR*); chromosomal insertion at *att*Tn7 site of *hsdR* sequence under control of its natural promoter (NOG899/902) | This study |
| PAO1 *PrecA-mscarlet* | PAO1 chromosomal insertion at *att*Tn7 site of P*recA-mscarlet* without antibiotic resistance cassette | This study |
| PAO1 Δ*hsdM PrecA-mscarlet* | PAO1 CRISPR-Cas9 deletion of *PA2735* (*hsdM*), chromosomal insertion at *att*Tn7 site of P*recA-mscarlet* without antibiotic resistance cassette | This study |
| ***Pseudomonas aeruginosa* clinical isolates** | | |
| PSAE1656 |  | Bactome database^44^ |
| F2034 |  | Bactome database^44^ |
| ZG5021922 |  | Bactome database^44^ |
| ZG205864 |  | Bactome database^44^ |
| ZG8525123 |  | Bactome database^44^ |
| F1880 |  | Bactome database^44^ |
| F1883 |  | Bactome database^44^ |
| MHH17783 |  | Bactome database^44^ |
| MHH0985 |  | Bactome database^44^ |
| MHH17767 |  | Bactome database^44^ |
| F1997 |  | Bactome database^44^ |
| CH5291 |  | Bactome database^44^ |
| PSAE2139 |  | Bactome database^44^ |
| F2005 |  | Bactome database^44^ |
| ESP075 |  | Bactome database^44^ |
| Pa16-13 |  | Weimann collection^41^ |
| F2003 |  | Bactome database^44^ |
| CF4S1(14) |  | Weimann collection^41^ |
| CF5S1(13) |  | Weimann collection^41^ |
| CF5S1(34) |  | Weimann collection^41^ |
| PSAE1656/pLOLA-RM type I |  | This study |
| PSAE1656/pLOLA-RM type II |  | This study |
| Pa16-13/pLOLA-RM type I |  | This study |
| Pa16-13/pLOLA-RM type II |  | This study |
| CF4S1(14)/pLOLA-RM type I |  | This study |
| CF4S1(14)/pLOLA-RM type II |  | This study |
| CF5S1(13)/pLOLA-RM type I |  | This study |
| CF5S1(13)/pLOLA-RM type II |  | This study |
| CF5S1(34) )/pLOLA-RM type I |  | This study |
| CF5S1(34) )/pLOLA-RM type II |  | This study |
| ***Escherichia coli*** | | |
| SM10λpir | *thi thr leu tonA lacY supE recA*::RP4-2-TcR::Mu KmR λpir | ^76^ |
| **Plasmids** | | |
| pS448•CsR | oriV (pRO1600/ColE1), *xylS* (devoid of internal Eco31I restriction site), P_m_→*cas9*, P_EM7_→sgRNA, Sm^R^ |  |
| pS448•CsR_*hsdM* | oriV (pRO1600/ColE1), *xylS* (devoid of internal Eco31I restriction site), P_m_→*cas9*, P_EM7_→sgRNA(*hsdM*), Sm^R^ |  |
| pS448•CsR_*hsdR* | oriV (pRO1600/ColE1), *xylS* (devoid of internal Eco31I restriction site), P_m_→*cas9*, P_EM7_→sgRNA(*hsdR*), Sm^R^ |  |
| pSH124-*ssr* | oriV RK2 lacI^q^,P_trc_→*ssr*, Amp/Carb^R^ |  |
| pUC18T-mini-Tn7T-Gm | Gm^r^ on mini-Tn*7*; for gene insertion in Gm^s^ bacteria | ^61^ |
| pUC18T-mini-Tn7T-Gm P*_hsdM_*-*hsdM* | Gm^r^ on mini-Tn*7*; for gene insertion in Gm^s^ bacteria. Sequence of P*_hsdM_*-*hsdM* | This study |
| pUC18T-mini-Tn7T-Gm P*_hsdR_*-*hsdR* | Gm^r^ on mini-Tn*7*; for gene insertion in Gm^s^ bacteria. Fusion of P*_hsdR_*-*hsdR* |  |
| pTNS2 | R6K γ ori (*tnsABCD*) | ^61^ |
| pFLP2 | Ap^R^ Cm^R^ r*epA*(Ts); pSC101-based vector expressing the Flp recombinase | ^61^ |
| pLOLA |  | ^31^ |

**Extended Data Table 5** Strains and Plasmids used in this study.
