## Extended Table 6 for "Loss of restriction-modification methyltransferases drives persistence to fluoroquinolones in *Pseudomonas aeruginosa*"

| **Primer Name** | | **Primer Sequence (5´- 3´)** |
| --- | --- | --- |
| **CRISPR/Cas9 spacer and sequencing primers** | | |
| NOG844 (*Sp_hsdM_F*) | | GCGCGGTCTTCCAACTCTTGCCGTA |
| NOG845 (*Sp_hsdM_R*) | | AAACTACGGCAAGAGTTGGAAGACC |
| NOG855 | | TCTGGCTCATGACCTGCTTG |
| NOG856 | | AACGCCCAGCATTTGGTTTC |
| NOG862 (*Sp_hsdR_F*) | | GCGCGCCGGGGGCCATCTATGCACT |
| NOG863 (*Sp_hsdR_R*) | | AAACAGTGCATAGATGGCCCCCGGC |
| NOG864 | | ACGAAATGGCGCAACTTTCC |
| NOG865 | | GCTAGGAACGCGGATAGCTT |
| **CRISPR/Cas9 recombineering oligonucleotides** | | |
| NOG857 | | GTGGCTTGCCACTAAACATATTTGAATGAAGAGTTGAAGCGCTGATAATGTGATGACAGCCACGAGCAGATATCCCAGCTACCGTGTATCCGGCCTCCCT |
| NOG857 | | GACAATAAGCCGGGCAAGAAGAAGGGAGACGACGCATGAAACCCACCGATTAGTCTGAAAACACTGTTGGGCTGGTTTCCAGTGCTGCGAAAATTATTCC |
| **Routine primer for checking the presence of *ssr*- and *cas9*-carrying plasmids** | | |
| AAR78 (ssr_F) | | TGAGCCAAGTAGCCAGGGTCG |
| AAR79 (ssr_R) | | GATTTGTAGACGTTGGCAGCGCAG |
| AAR80 (cas9_F) | | AAGCACAAGTGTCTGGACAAGG |
| AAR81 (cas9_R) | | TCACTCAAACCTCCACGTTCAGC |
| **Chromosomal complementation** | | |
| NOG897 | ATGC**GAGCTC**TTCCTTTTGTGAAAGTACTTGTGATCGAC | |
| NOG898 | GCAT**GGTACC**TCAAGCTGTCCCCACGATCTT | |
| NOG899 | ATGC**GAGCTC**AGCGGCGGAATACCTGACGT | |
| NOG900 | GGTTTCATGCGTCGTCTCCCTTGTCGTAGTCGATGGACGTGGC | |
| NOG901 | GCCACGTCCATCGACTACGACAAGGGAGACGACGCATGAAACC | |
| NOG902 | GCAT**GGTACC**CTATCGCCCATGCGCAATCCT | |

**Extended Data Table 6** Primers used in this study
