## Supplementary figures and images for "Loss of restriction-modification methyltransferases drives persistence to fluoroquinolones in *Pseudomonas aeruginosa*"

### Extended Figure 1

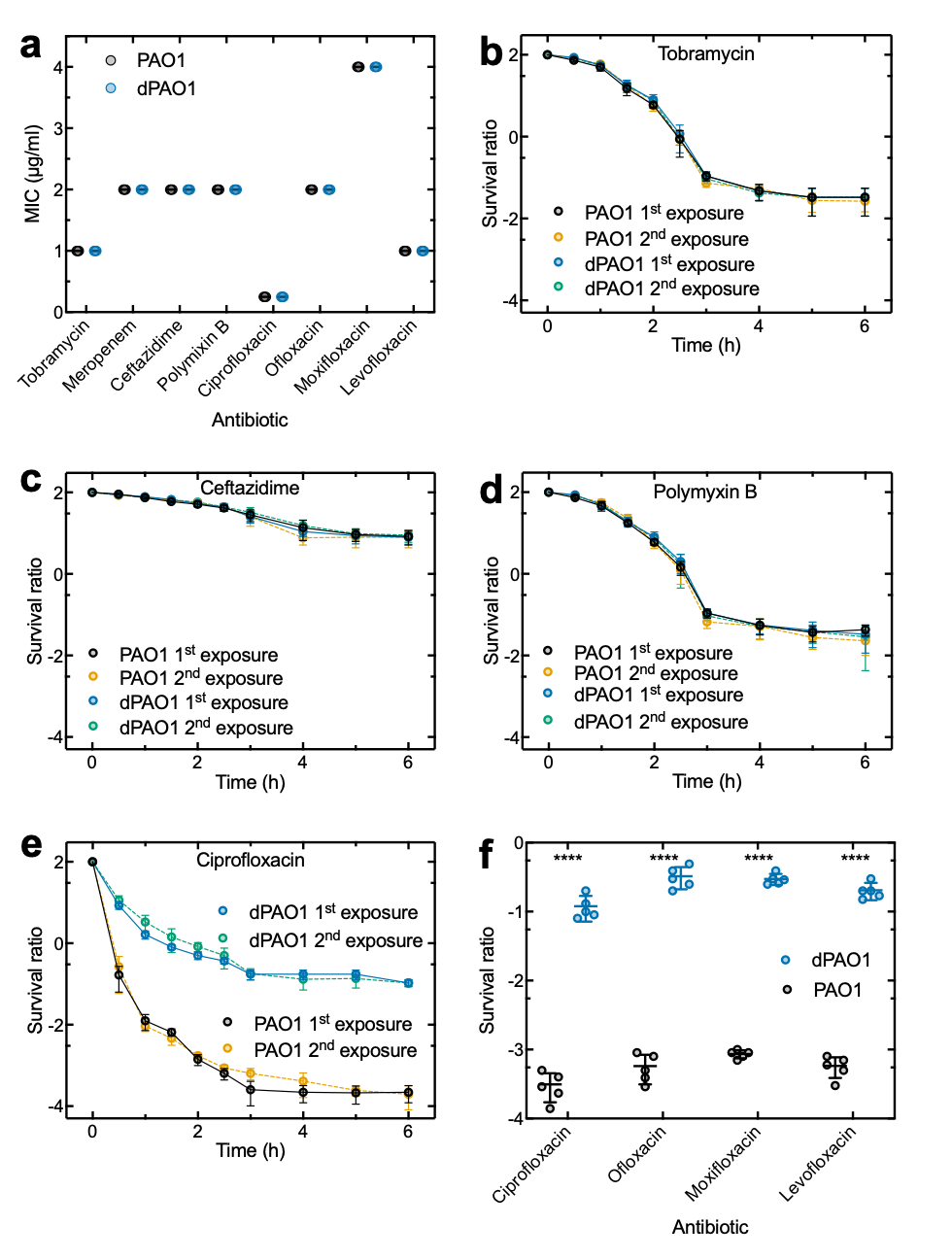

### Extended Figure 2

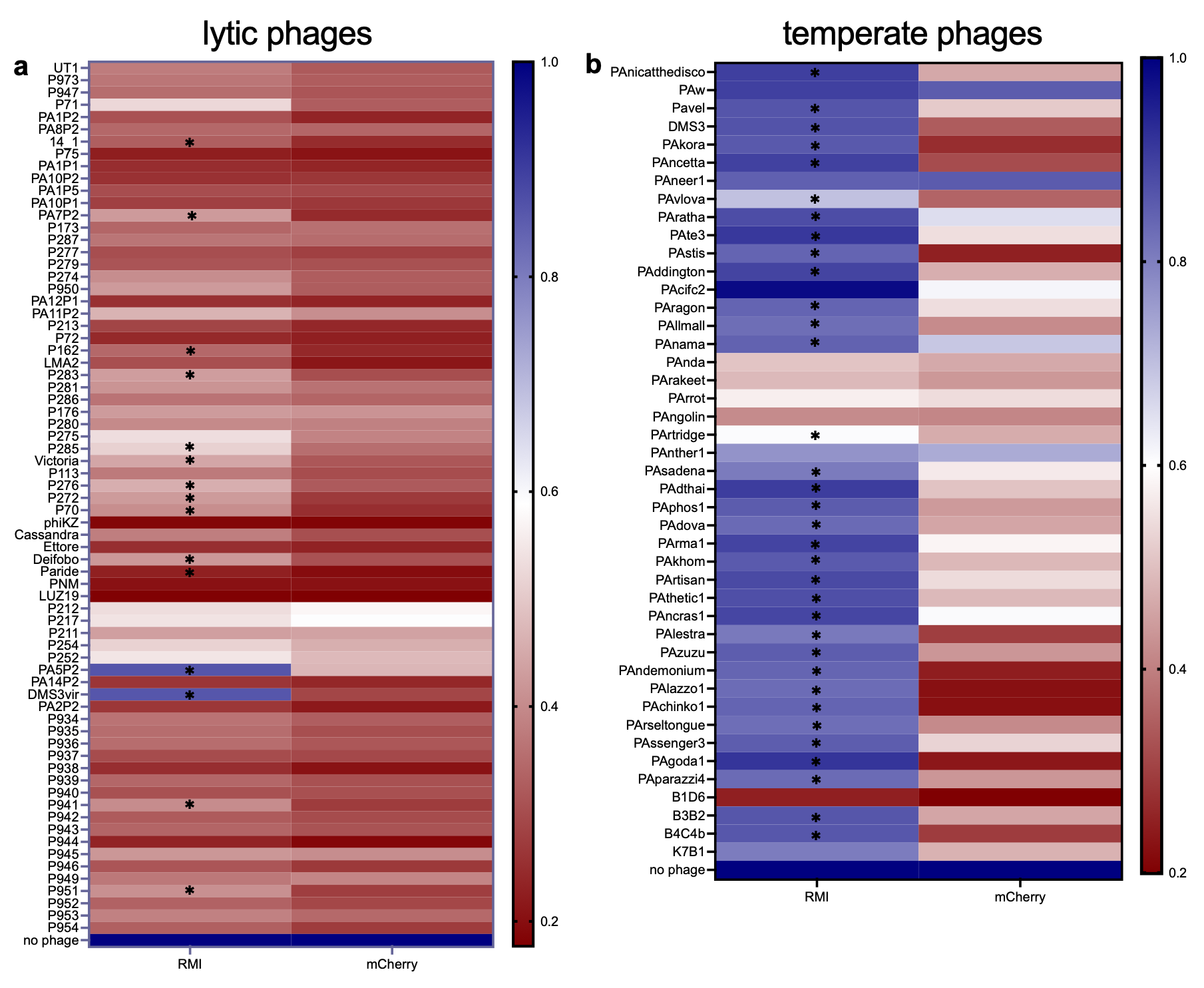

### Extended Figure 3

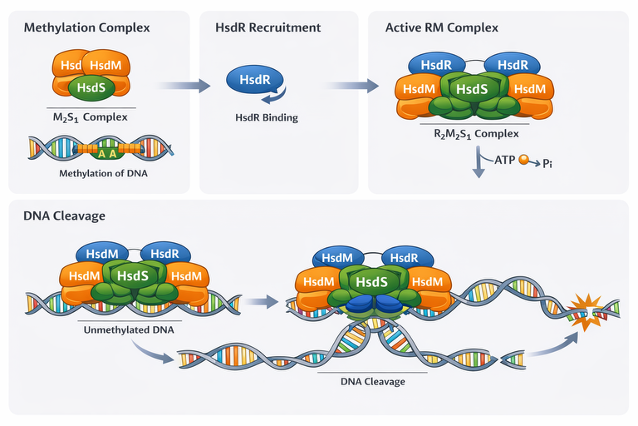

### Extended Figure 4

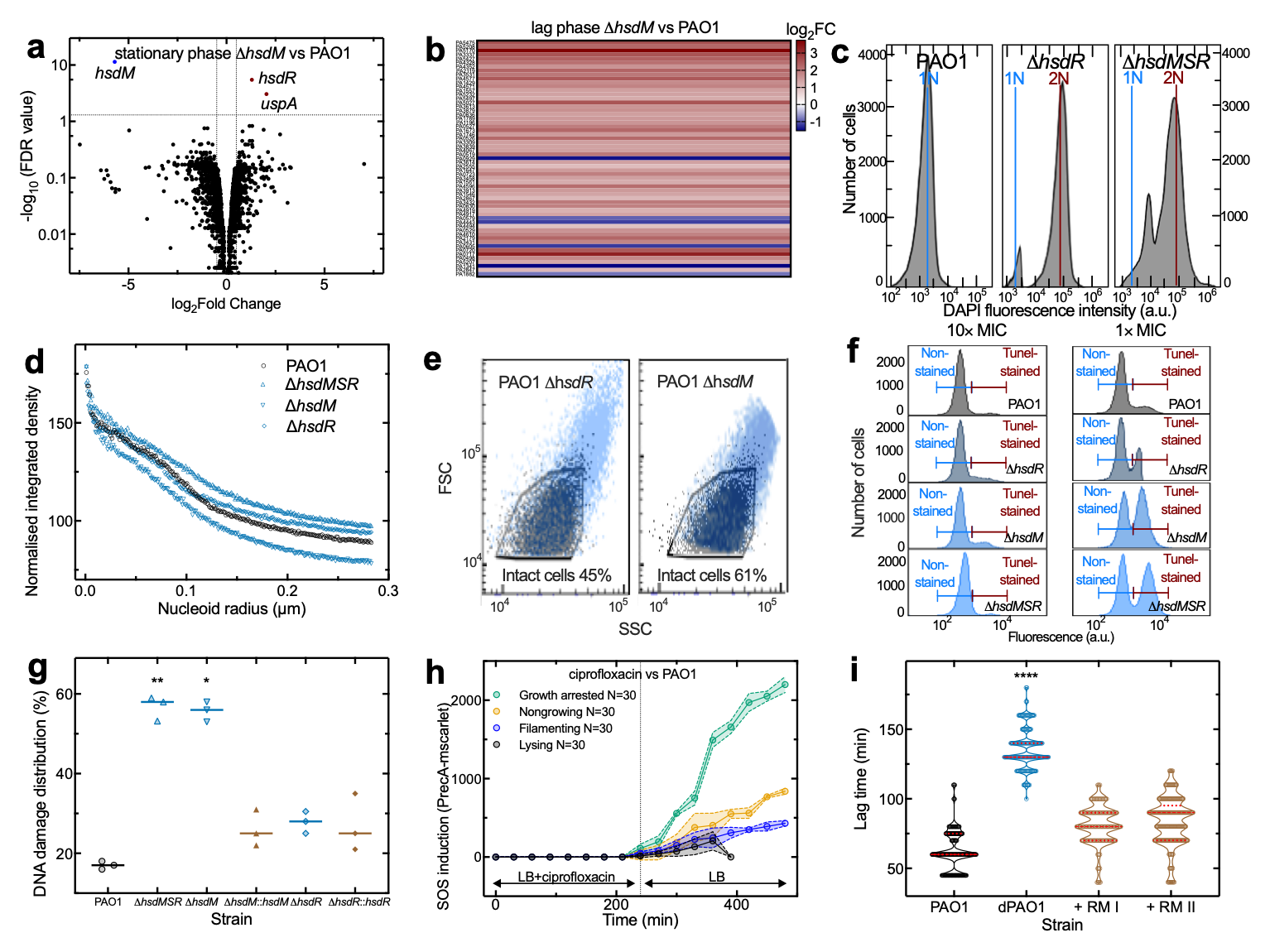

### Extended Figure 5

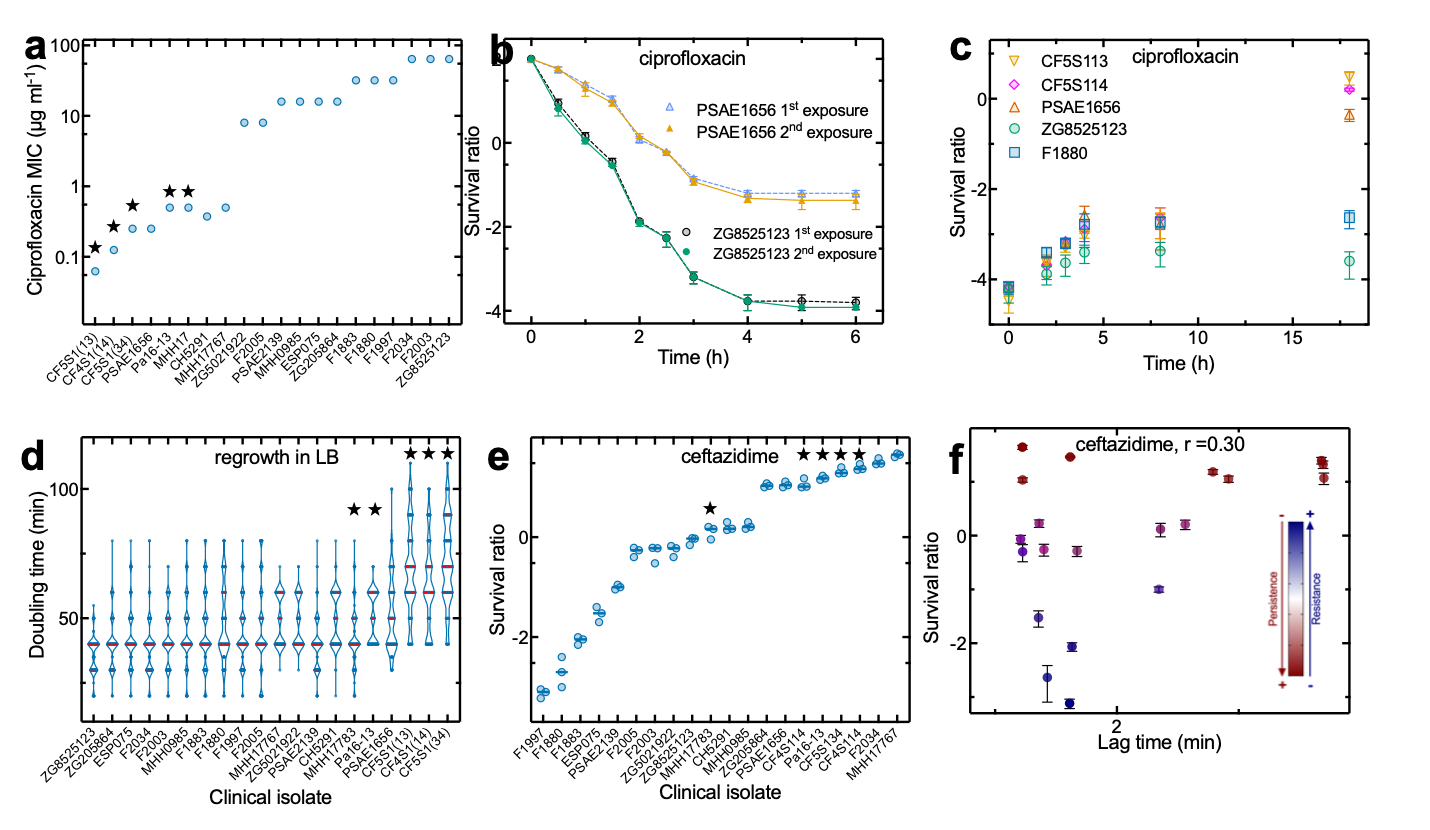

### Extended Figure 6

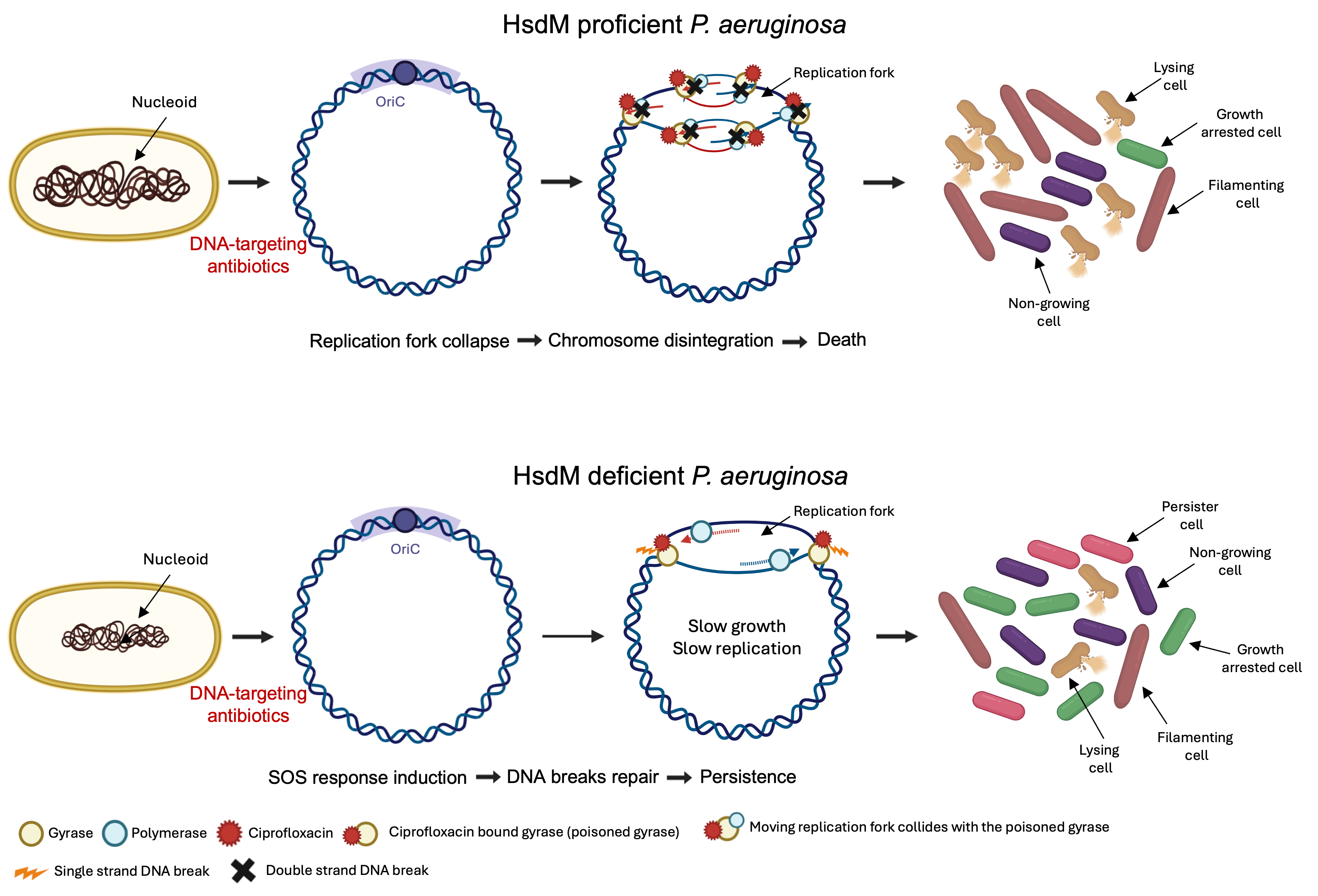
